# A personalizable autonomous neural mass model of epileptic seizures

**DOI:** 10.1101/2021.12.24.474090

**Authors:** Edmundo Lopez-Sola, Roser Sanchez-Todo, Èlia Lleal, Elif Köksal-Ersöz, Maxime Yochum, Julia Makhalova, Borja Mercadal, Maria Guasch, Ricardo Salvador, Diego Lozano-Soldevilla, Julien Modolo, Fabrice Bartolomei, Fabrice Wendling, Pascal Benquet, Giulio Ruffini

**Affiliations:** Brain Modeling Department, Neuroelectrics Barcelona, Barcelona, Spain; Univ Rennes, INSERM, LTSI - UMR 1099, F-35000 Rennes, France; Assistance Publique – Hôpitaux de Marseille, Service d’Epileptologie et de Rythmologie Cerebrale, Hopital La Timone, Marseille, France; Aix-Marseille Université, Marseille, France

**Keywords:** Neural mass model, SEEG Recordings, depolarizing GABA, epilepsy

## Abstract

Work in the last two decades has shown that neural mass models (NMM) can realistically reproduce and explain epileptic seizure transitions as recorded by electrophysiological methods (EEG, SEEG). In previous work, advances were achieved by increasing excitation and heuristically varying network inhibitory coupling parameters in the models. Based on these early studies, we provide a laminar NMM capable of realistically reproducing the electrical activity recorded by SEEG in the epileptogenic zone during interictal to ictal states. With the exception of the external noise input into the pyramidal cell population, the model dynamics are autonomous. By setting the system at a point close to bifurcation, seizure-like transitions are generated, including pre-ictal spikes, low voltage fast activity, and ictal rhythmic activity. A novel element in the model is a physiologically motivated algorithm for chloride dynamics: the gain of GABAergic post-synaptic potentials is modulated by the pathological accumulation of chloride in pyramidal cells due to high inhibitory input and/or dysfunctional chloride transport. In addition, in order to simulate SEEG signals for comparison with real seizure recordings, the NMM is embedded first in a layered model of the neocortex and then in a realistic physical model. We compare modeling results with data from four epilepsy patient cases. By including key pathophysiological mechanisms, the proposed framework captures succinctly the electrophysiological phenomenology observed in ictal states, paving the way for robust personalization methods based on NMMs.

## 1. Introduction

Computational models have proven to be a powerful tool to understand the pathophysiology underlying epileptic activity. In [1], neural mass models (NMM) were used to represent realistic epileptic seizure transitions: changes from interictal to ictal state were achieved by increasing the excitatory synaptic gain at the level of glutamatergic pyramidal cells and varying the inhibitory synaptic gains of GABAergic interneurons. This model simulated the transition through some of the phases typically observed in epileptic seizures recorded with stereoelectroencephalography (SEEG), such as pre-ictal spikes, low voltage fast onset activity and ictal rhythmic activity.

The NMM framework describes the dynamics of the average membrane potential and firing rates of populations of neurons in a cortical column [2, 3, 4, 5]. A NMM consists of second order differential equations describing the membrane perturbation of a neuronal population at the level of each synapse due to the input currents from another neuronal population, plus an equation converting membrane potential into firing rate using Freeman’s sigmoid function [6]. Models of several populations can be coupled to produce different types of activity (see [7] for a complete description of the Jansen and Rit three population model), including epileptiform activity [1].

There is increasing evidence of the role of GABAergic transmission in seizure initiation, propagation and termination, both in the immature and mature brain (see [8] for an extensive review). GABAergic synaptic transmission can have seizure-promoting effects when release of GABA neurotransmitters leads to excitation instead of inhibition, a phenomenon known as “depolarizing GABA”. Indeed, optogenetic control of perisomatic-targeting GABAergic interneurons such as parvalbumin-positive (PV) cells showed that high frequency discharge of this cell types was able to trigger fast onset activity [9, 10], and selective activation of PV cells and another sub-class of GABAergic interneurons, somatostatin-positive (SST) cells, triggered epileptiform activity [11].

Mechanistically, the dysregulation of chloride homeostasis observed in epileptic tissues appears to play a major role in the depolarizing effect of GABA_A_ neurotransmitters during epileptic seizures [12], and there is evidence for the accumulation of chloride in pyramidal cells at seizure onset [13]. Changes in the expression level of chloride cotransporters, such as KCC2 (K-Cl cotransporter type 2) and NKCC1 (Na-K-2Cl cotransporter type 1) have been identified as the main causes for pathological chloride accumulation in the epileptic tissue [8, 14, 15, 16, 17].

Recently, a computational model including activity-dependent GABA depolarization was used to reproduce epileptic discharges in Dravet syndrome patients [18]. In the model, GABA depolarization was mediated by PV-cells and was represented by a parameter that was gradually increased to generate transitions from background to interictal, low voltage fast-onset seizure initiation, ictal-like activity and seizure-like activity termination.

Building on previous work [1, 18] and on data reported in the literature [8], we have extended the NMM formalism to include chloride dynamics. By modeling pathological chloride accumulation in the pyramidal cell population, we can simulate autonomous, realistic epileptic seizures. Interestingly, the model accurately reproduces the sequence of complex epileptiform patterns observed during the interictal to ictal transition. The model is autonomous in the sense that it is directly driven to ictal state by random fluctuations of external stochastic input onto the pyramidal cells (which represent fluctuations of cortico-cortical or thalamo-cortical input, or neuromodulatory effects).

Developing personalized computational models that simulate brain activity during epileptic seizures is of crucial importance for a better understanding and optimization of treatments. Not only can models help identifying the pathophysiological mechanisms involved in seizure initiation, propagation and termination, but also they can improve the efficacy of potential treatments that range from the surgical resection of epileptogenic tissue [19, 20, 21] to non-invasive brain stimulation. In this regard, model parameters should be fine-tuned to fit the main patient-specific features of the SEEG recordings.

The generation of realistic SEEG data to compare with real recordings requires combining the NMM framework with a physical head model that accounts for both the location of the SEEG electrode contacts and the biophysical features of the volume conductor (tissue morphology and conductivity). This combination is crucial to solve the SEEG-forward problem and hence simulate signals analogous to real SEEG signals. The present model includes a laminar architecture that represents the cortical layers of the human brain, as was done in previous work [22, 23]. We can then embed the seizure-generating NMM into a physical head model to generate SEEG-like activity during the transition from interictal to ictal state. We provide here examples of model personalization, where we compare the simulated SEEG signal with the recordings of four epilepsy patients during seizures. We also present a proof of concept of stimulation with constant weak electric fields for the reduction of seizure activity in the model.

## 2. Methods

### 2.1. Patient selection and data collection

The four patients included in this study required SEEG as part of usual clinical care. Clinical information about the patients can be found in Table 1. In all cases, several intracerebral electrodes were placed in different brain regions of the patient based on the clinician’s hypotheses about the localization of the epileptogenic zone. For each patient, several seizures were recorded by the SEEG electrodes: three seizures for Patients 1 and 2, four seizures for Patient 3 and two seizures for Patient 4. All seizure data was included in the analysis.

**Table 1:** Clinical characteristics of the patients included in the study.

| Patient ID | 1 | 2 | 3 | 4 |
| --- | --- | --- | --- | --- |
| Gender | F | F | F | M |
| Age at epilepsy onset (y) | 14 | 9 | 10 | 11 |
| Epilepsy duration (y) | 7 | 14 | 14 | 4 |
| Epilepsy type | Parietal | Occipital | Temporal | Temporal |
| Surgical procedure | SR | SR | TC, SR | TC, SR |
| Surgical outcome (Engel class) | I | II | I | I |
| MRI | R parietal DNET | Normal | R temporal FCD | Normal |
| Histopathology | DNET | FCD1 | FCD2 | FCD2 |
| Side | R | L | R | L |

Abbreviations: DNET, dysembryoplastic neuroepithelial tumor; FCD, focal cortical dysplasia; L, left; R, right; SR, surgical resection; TC, thermocoagulations.

### 2.2. SEEG recordings

SEEG recordings were obtained using intracerebral multiple contact electrodes with a 154-channel Deltamed system. Offline, SEEG data was high-pass filtered above 0.5 Hz. Bipolar derivations were taken by neighbour contact subtraction for each electrode.

For comparison with the simulated data, we have selected SEEG contacts presenting the most epileptogenic activity during seizure. The metric used for the identification of epileptogenic contacts was the Epileptogenicity Index (EI) [24]. In brief, the EI classifies SEEG contact signals according to the tonicity of rapid discharges and the delay with respect to seizure onset. For each patient, SEEG data from several seizure events was available. We computed the EI for each seizure using the open-source software AnyWave [25] and selected as most epileptogenic contact the contact with the highest average EI over all seizures.

The location of the most epileptogenic contacts in each patient is described hereafter:

- *Patient 1* : right supramarginal gyrus, directly inside an anatomical lesion suspected to be the epileptic focus;
- *Patient 2* : left occipital pole, no anatomical lesion was known for this patient;
- *Patient 3* : right posterior superior temporal sulcus, in a region of cortical dysplasia suspected to be the epileptic focus;
- *Patient 4* : left hippocampus and left posterior T2 (middle temporal gyrus), with no specific lesion for this patient.

A 3D view of the electrode contacts position in the patients’ head model can be seen in Figure S1.

### 2.3. SEEG data analysis

Seizures were identified by a trained neurologist who set the time of seizure onset. In order to optimize the trade-off between time and frequency resolution, we carried out separate time-frequency representations (TFRs) of oscillatory power for slow and fast frequencies [26]. The TFRs allowed us to easily identify canonical features of each seizure: a fast onset characterized by low-amplitude high frequency chirp-like bursts and a rhythmic clonic phase characterized by large amplitude sustained oscillations. For each patient and seizure, we manually defined the onset and offset of the two events and estimated their duration and peak frequency. The peak frequency was obtained by spectral parametrization following the approach described in [27] (more details are given in Appendix A).

Figure 1 shows examples of SEEG seizure recordings for each of the four patients. For patients 1, 2 and 3, all seizures displayed a phase of low-voltage fast activity (LVFA) at seizure onset followed by rhythmic oscillations. The frequency of the LVFA period was much higher for patients 1 and 3 (∼100 Hz) than for patient 2 (∼40 Hz). Pre-ictal spikes were recorded prior to the LVFA phase for patients 2 and 3. For Patient 4, a burst of spikes in the beta range was followed by rhythmic oscillations in all the seizures recorded. The SEEG seizure recordings for all seizures and all patients are shown in Figure S2.

**Figure 1:**
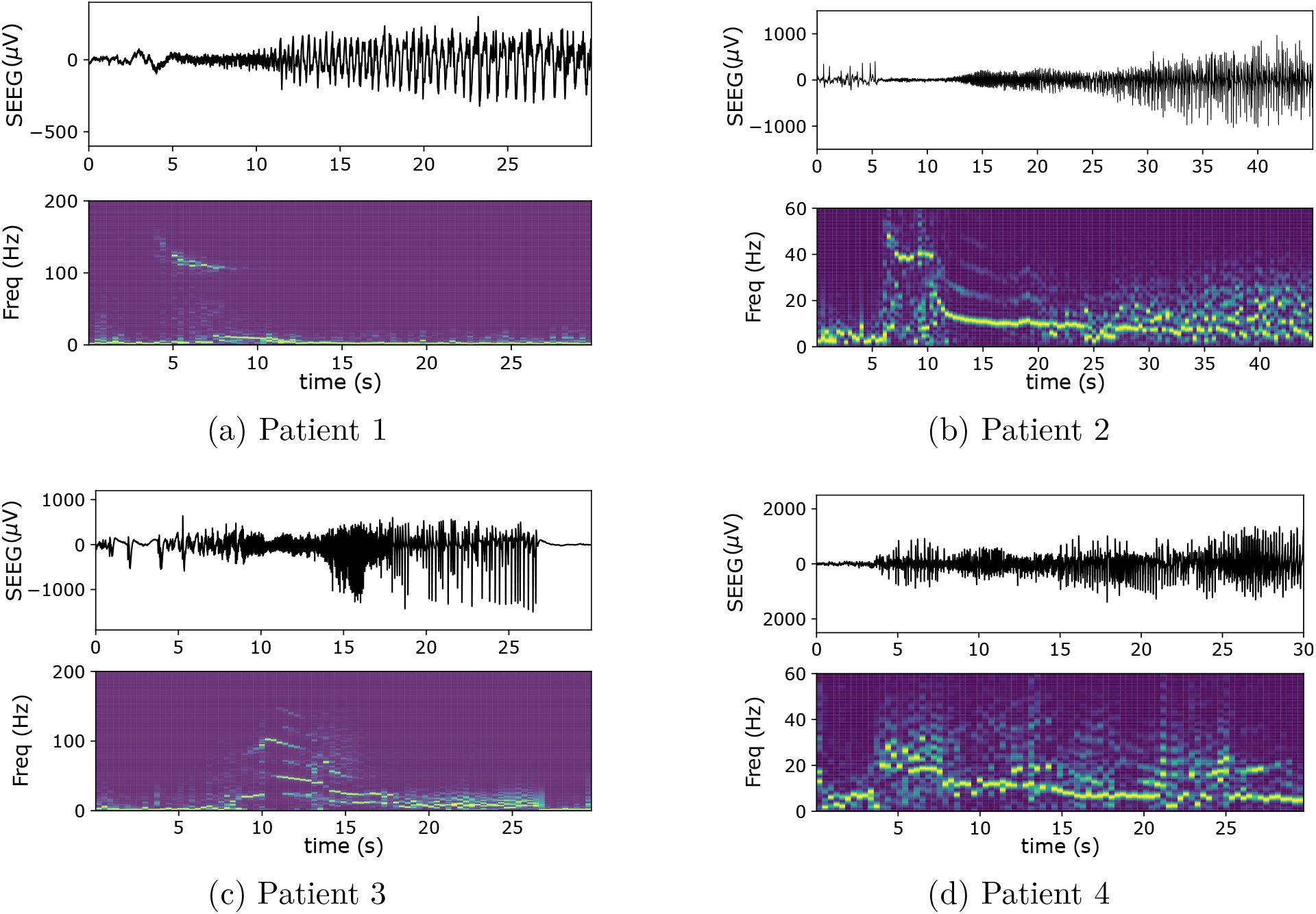
Examples of SEEG seizure recordings for each of the four patients. Low-voltage fast activity (LVFA) was present at seizure onset for patients 1, 2 and 3. Preictal spikes were recorded prior to the LVFA phase for patients 2 and 3. For Patient 4, seizure onset was characterized by a burst of spikes in the beta range.

### 2.4. Laminar NMM with dynamic inhibitory synaptic gains

We have recently extended the NMM framework to be able to represent the physics of synaptic electrical current flow in cortical layers [22, 23]. The model developed here is such a *laminar NMM* consisting of populations of (i) pyramidal neurons (P), (ii) other excitatory cells (E), which represent lateral excitation by spiny stellate cells or other pyramidal neurons in layer IV, (iii) fast inhibitory interneurons (PV), such as parvalbumin-positive cells releasing GABA_A_ and targeting the soma and proximal dendrites of pyramidal cells, and (iv) slow inhibitory interneurons (SST), such as somatostatin-positive cells targeting the pyramidal cells apical dendrites releasing GABA_A_.

With regard to connectivity, the pyramidal population sends excitatory input to all other neuronal populations and receives excitatory signals from the excitatory population and inhibitory signals from the fast and slow inhibitory interneurons. The NMM also includes an external excitatory input to the main pyramidal population, which represents the input from other cortical columns or subcortical structures. Based on the extensive literature describing local connections in cortical columns, we have also included in the model autaptic self-connections in the fast inhibitory interneuron population, and synaptic connections from slow inhibitory interneurons to fast inhibitory interneurons [28]. The resulting circuit architecture is a slight variation of the models proposed in [1] and [18]. The laminar locations of each neuronal population and their synapses are derived from the available literature [28, 29]. Figure 2 summarizes the complete architecture used in the present work.

**Figure 2:**
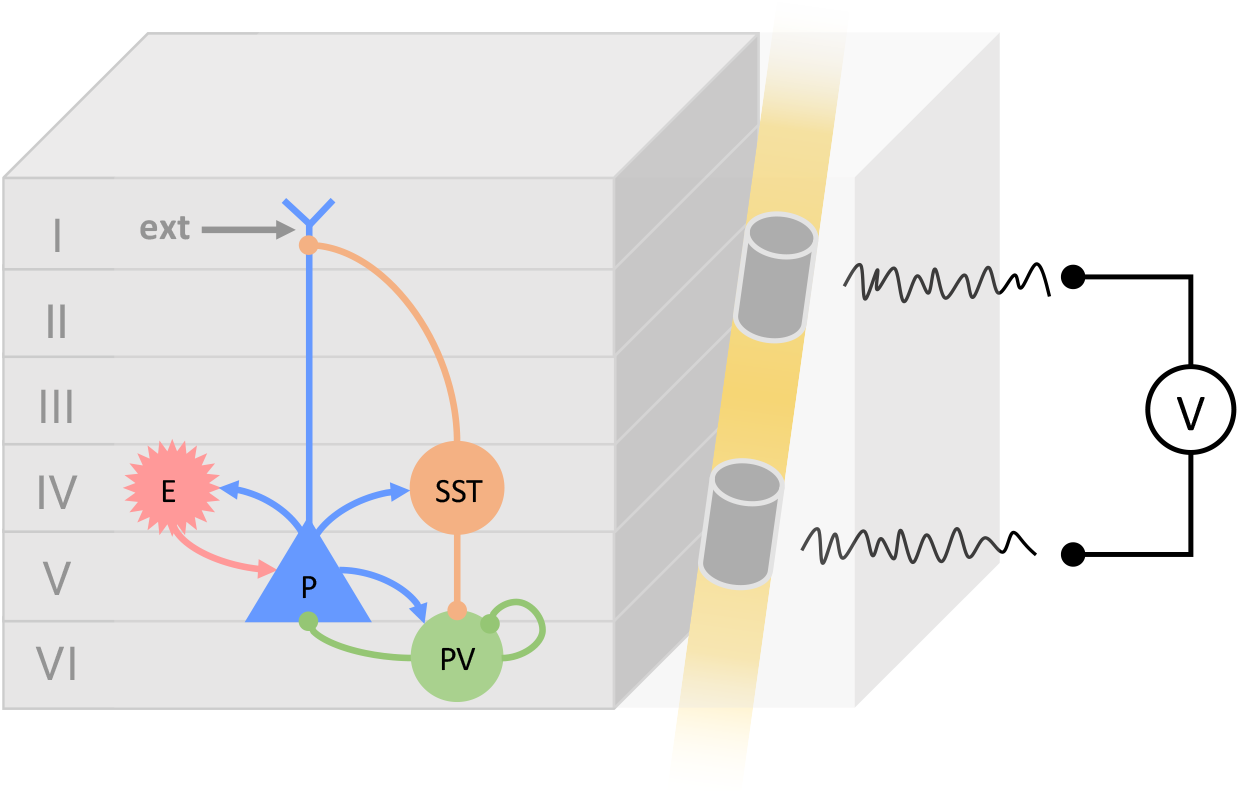
Laminar neural mass model architecture, including four different neuronal populations: pyramidal cells (P), other excitatory cells (E), dendritic projecting interneurons with slow synaptic kinetics (SST) and somatic-projecting interneurons with faster synaptic kinetics (PV). An external input to the pyramidal cell population from other cortical areas and subcortical structures is included (*ext*). Arrows represent excitatory synapses and dot terminals represent inhibitory synapses. Neurons (which are now extended objects spanning several layers) and synapses are placed in specific cortical layers, a key aspect for the computation of realistic SEEG signals. A physical model is included to simulate the bipolar voltage measured by the SEEG electrode contacts inside (or close to) the active patch.

We note that the proposed circuit is meant to be representative of the architecture found in an average cortical area, although we acknowledge that the neuronal populations found in some regions will differ in type, connectivity and distribution. This might be especially true for damaged cortical structures, as is the case for some of the patients included in the study (see Table 1). We have decided for simplicity to use the same cortical architecture for all patients and to then include some parameter modifications to represent the pathological character of the epileptic tissue, such as the synaptic gain of excitatory connections or the connectivity strength of synapses (see Section 2.6).

We present a synapse-driven formalism, where each synapse *s* is described by an equation representing the conversion from an input firing rate *φ*_*n*_ from the pre-synaptic population *n* into an alteration of the membrane potential *u*_*s*_ of the post-synaptic neuron. This relation is represented by the integral operator 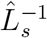 (a linear temporal filter), the inverse of which is a differential operator 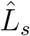,

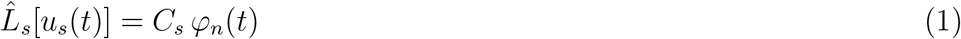

where *C*_*s*_ is the connectivity constant between the populations and the differential operator 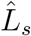 is defined as

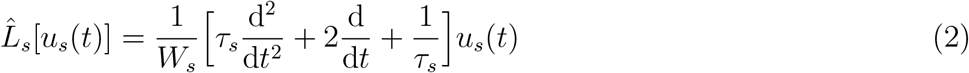

with *W*_*s*_ the average excitatory/inhibitory synaptic gain and *τ*_*s*_ the synaptic time constant. In classical neural mass models, *W*_*s*_ is a constant parameter associated with each synapse type [1, 30]. Here, to account for the chloride-dependent variations of the PSP amplitude generated by GABA release, the gain *W*_*s*_ of some of the synapses (SST → P and PV → P) is a dynamic variable *W*_*s*_(*t*) governed by chloride transport and other equations (see next section).

A neuronal population *𝒫*_*n*_ state is characterized by its membrane potential *v*_*n*_ (the summation of all its pre-synaptic membrane perturbations *u*_*s*_) and by its firing rate *φ*_*n*_, which is computed using a non-linear function of the membrane potential,

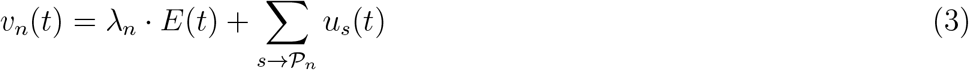

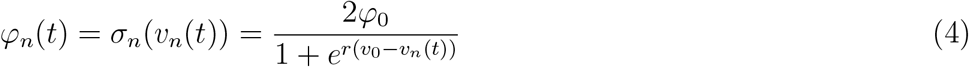

where the sum in the first equation is over synapses reaching the population, *φ*_0_ is half of the maximum firing rate of each neuronal population, *v*_0_ is the value of the potential when the firing rate is *φ*_0_ and *r* determines the slope of the sigmoid at the central symmetry point (*v*_0_, *φ*_0_) [6, 5, 7].

The term *λ*_*n*_ · *E*(*t*) represents the membrane perturbation induced by an external electric field [31, 32, 33] and accounts for the effects of electrical stimulation or ephaptic effects [34], in the case where they are to be included (see Section 3.4). The *λ · E* model assumes that the electric field effect is proportional to the projection of the electric field on the axon direction. Since the electric field generated by tES is thought to affect mainly elongated neurons such as pyramidal cells [31], the *λ*_*n*_ · *E*(*t*) term affects only the membrane potential of the pyramidal population in our model.

The detailed equations for each synapse in the model and a description of the model parameters are provided in Appendix B. An illustrative diagram of the model equations is shown in Figure 3.

**Figure 3:**
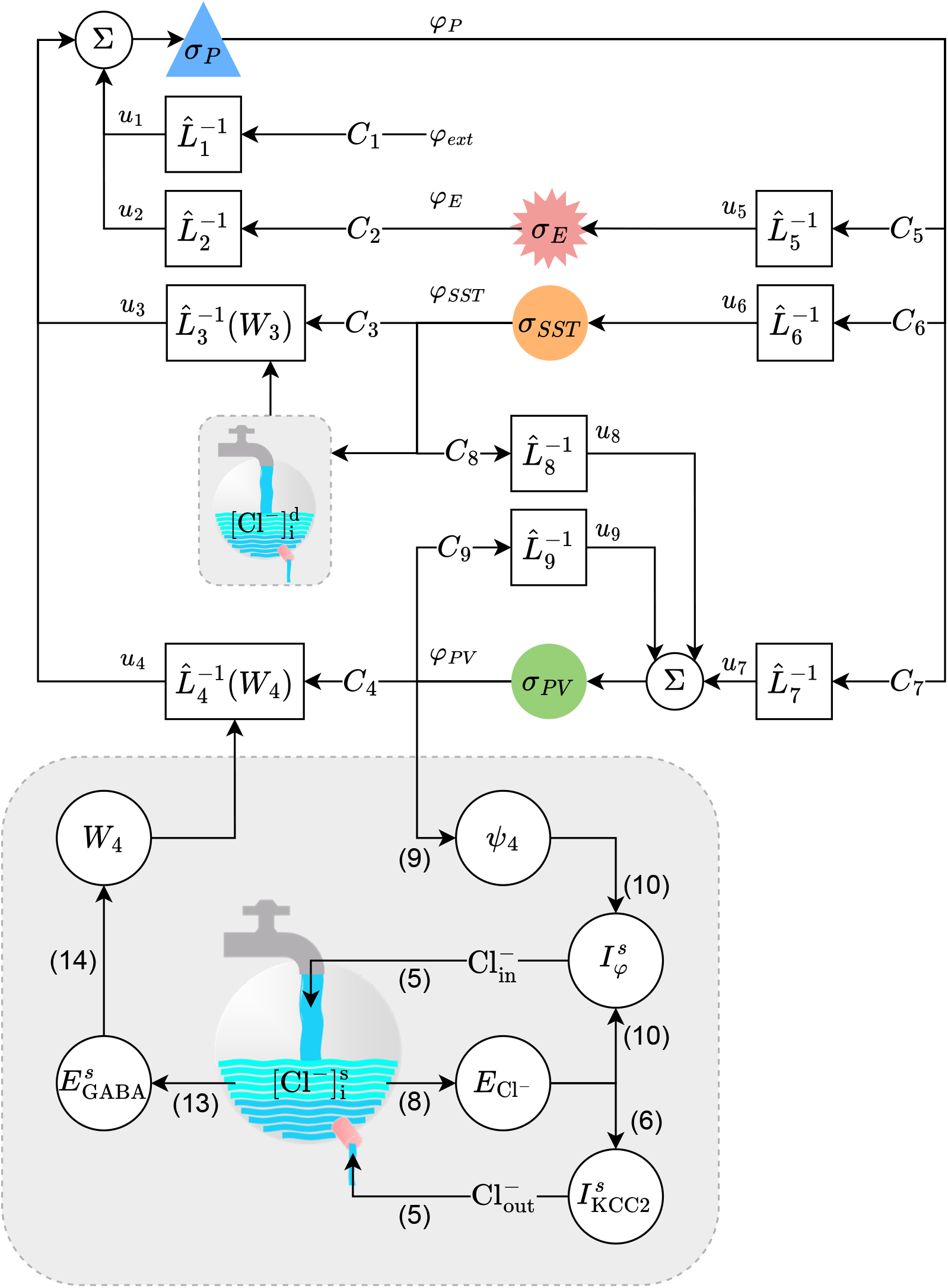
Diagram of the extended NMM equations. The grey box below represents the equations for chloride dynamics of the PV→P synapse (chloride dynamics in the soma, superscript *s*). Equation numbers corresponding to the main text are displayed in parenthesis. The model for chloride accumulation dynamics uses as input from the standard NMM equations the firing rate of the inhibitory interneurons (*φ*_*PV*_ in this case), and update the synaptic gain of the connection (*W*_4_). The grey box in the middle of the diagram, marked with 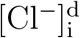, represents the equations for chloride dynamics equations of the SST→P synapse (chloride dynamics in the dendrites), which are described by the same model as the one of the bottom grey box.

### 2.5. Model for chloride accumulation dynamics

To the synapse equations presented in the previous section we add supplementary ODEs to account for the dynamics of Cl^−^ accumulation in the pyramidal cell population and its impact on synaptic gains. In the model, the dynamics of Cl^−^ accumulation in the pyramidal cell population are driven by the release of GABAergic neurotransmitters from the fast and slow inhibitory interneurons [12]. Given that those interneurons target two distinct cell locations (PV cells target more frequently perisomatic locations, while SST cells usually target the apical dendrites [28]), we modeled the Cl^−^ dynamics separately for each synaptic type location.

As in previous work [35, 36], the concentration of Cl^−^ in a synaptic location is calculated according to the balance equations as a function of the combined extrusion of chloride by KCC2 transporters and the firing-rate-dependent influx of chloride in GABA_A_ synapses, modulated by a surface-to-volume and charge-to-concentration translating parameter 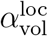 that depends on the synapse location (apical dendrites or perisomatic region),

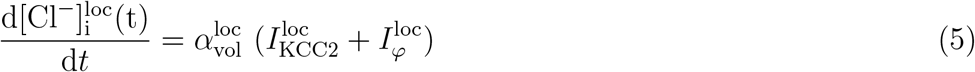

The parameter 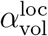 reflects, at the population level, the location-dependent modulation of the impact of ionic currents on the change of Cl^−^ accumulation by the local cell compartment volume [35, 37]: the same ionic current will cause different Cl^−^ accumulation changes depending of the size of the reservoir, which depends on the synapse location in the cell. Due to the volume differences of apical dendrites compared to the soma, chloride influx or extrusion causes faster changes in the total Cl^−^ concentration in apical dendrites, as suggested by recent studies [12, 8]. Thus, 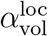 will be higher in the apical dendrites. We have assumed for simplicity that the chloride concentration in the soma and dendrites is independent, and that there is no diffusion of [Cl^−^] between them.

The Cl^−^ current associated with KCC2 transporters is given by

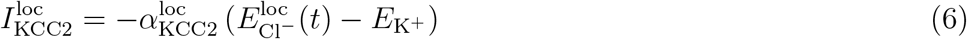

We assume here that the reversal potential of potassium 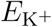 remains constant. The parameter *α*_KCC2_ reflects the rate of extrusion of Cl^−^ through KCC2 transporters. Since KCC2 transporter dysfunction plays a crucial role in pathological Cl^−^ accumulation in epilepsy [8, 12], *α*_KCC2_ is personalized to reflect patient epileptogenicity levels in different nodes: the most epileptogenic areas will display reduced KCC2 transporter function (low *α*_KCC2_), while healthy regions will have nominally functional KCC2 transporters (high *α*_KCC2_). Indeed, for a given set of parameters that produces seizure-like behavior, increasing *α*_KCC2_ is sufficient to avoid the transition to ictal state (see Figure E1). Some studies suggest that NKCC1 dysfunction might also contribute to the pathological accumulation of Cl^−^ in epilepsy [38]. As opposed to KCC2 transporters, NKCC1 transports Cl^−^ inside the cell. The model used here does not explicitly display the contribution of NKCC1 transporters Cl^−^ homeostasis. Given the opposite role of both transporters, the current model can be easily generalized to account for a balance of in/out transport of Cl^−^ by KCC2 and NKCC1 transporters without the need to modify substantially the current implementation.

The Cl^−^ current through GABA_A_ channels due to the activity of inhibitory interneurons is given by

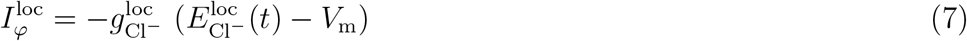

where 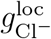 is the chloride conductance through GABAergic channels in a given location of the cell, *V*_m_ is the membrane potential of the neuron and 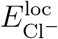 is the reversal potential of chloride in the cell location. For simplicity, for the purpose of calculating these currents, we have also assumed that the membrane potential of the cell *V*_m_ is not affected by Cl^−^ concentration dynamics. 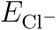 depends on the chloride accumulation inside the cell, and is given by Nernst-Planck equation,

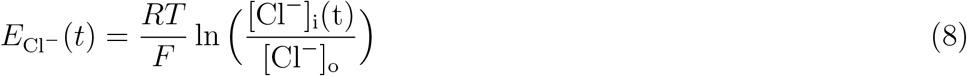

In the model, the influx of Cl^−^ into the cell is due to the release of GABA by inhibitory interneurons. Therefore, the conductance 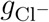 is proportional to the firing rate of the pre-synaptic GABAergic interneurons,

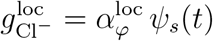

where *ψ*_*s*_(*t*) is a filtered version of the firing rate, which describes the average net synaptic flux for a given pre-synaptic firing rate *φ*_*n*_. The expression of *ψ*_*s*_(*t*) is based on exponential smoothing:

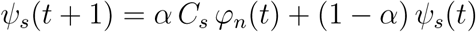

with *α* = 1 − *e*^Δ*t/τ*^ the smoothing factor. In differential form, we have:

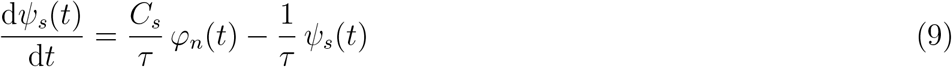

To compute the total input from inhibitory interneurons, the firing rate of each neuron is multiplied in Equation 9 by the number of connections from the pre-synaptic neuron to the pyramidal population *C*_*s*_. As before, the subscript *s* indexes the specific synapse involved (SST → P for the apical dendrites or PV → P for the soma) and *n* indicates the pre-synaptic population (SST or PV). The time constant of the exponential smoothing can be adapted to reflect current kinetics, which are slightly faster than the voltage kinetics. Here, for simplicity, we used the voltage time constants as well, i.e., we set *τ* = *τ*_*s*_. We can then express the chloride current through GABA_A_ channels due to the firing rate of inhibitory interneurons as

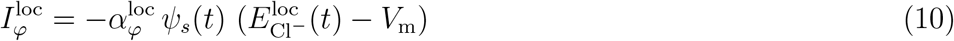

The parameter 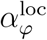 relates the activity of GABAergic interneurons with the Cl^−^ currents. This parameter can be personalized on a patient-specific basis.

The balance relation in Equation 5 is a differential equation for [Cl^−^] and requires an extra condition for its solution, i.e., the baseline value of [Cl^−^]. This value [Cl^−^]_0_ is thus another physiologically relevant model parameter leading to different system dynamics.

In the NMM framework, synaptic gains represent the population mean amplitude of the PSP generated by a given synapse type. The mean amplitude of the PSP generated by an ion species *A* in a synapse is proportional to the current generated by this ion species,

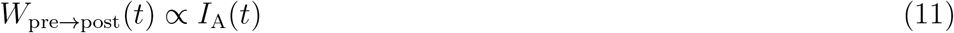

In the case of GABAergic synapses, the current is given by

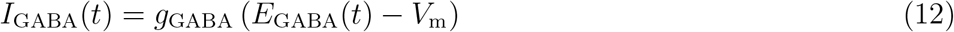

where *g*_GABA_ is the average inhibitory conductance, *V*_m_ is the membrane potential (assumed approximately constant) and *E*_GABA_ is the reversal potential of GABA (the joint reversal potential of Cl^−^ and 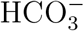), which can be calculated using the Goldman-Hodgkin-Katz equation,

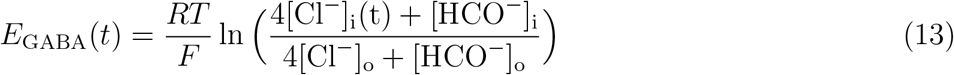

As mentioned above, we assumed that all ionic concentrations are constant to a good approximation except for the concentration of chloride inside the cells [Cl^−^]_i_. Following equations 11 and 12, the synaptic gains of inhibitory cell to pyramidal connections then are

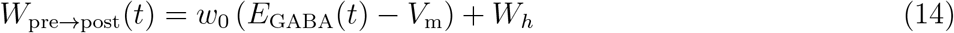

where *w*_0_ is the proportionality constant that links GABAergic currents to the mean amplitude of the PSP generated, and the term *W*_*h*_ represents the contribution of healthy cells to the average PSPs associated with a given population synapse. The rationale behind the additional term *W*_*h*_ is that there may be a proportion of healthy cells in the population where Cl^−^ will remain always within healthy levels, thus always generating non-pathological PSPs with a constant gain; this reflects the population-average nature of the analysis (see Appendix B.3 for more details).

The pathological chloride-dependent synaptic gains are in our model implemented in the connections from SST and PV populations to pyramidal populations, given by

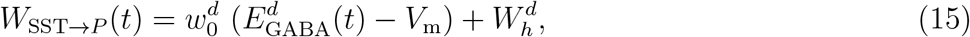

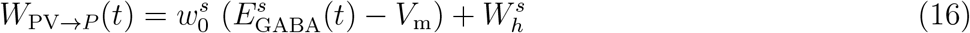

where the superscripts *s* and *d* indicate the synapse location (*s* for soma, *d* for dendrites). Other synaptic gains (pyramidal population to interneurons, SST to PV, etc.) are assumed to be constant in the model. The modeling choice of associating chloride dynamics to the synaptic gains of these two specific connections is based on the fact that it is sufficient to alter these two parameters to achieve realistic transitions from interictal to ictal state, as demonstrated in [1]. Future work should study whether including pathological chloride accumulation in other neuronal populations (PV cells due to the autaptic PV connection, for example) leads to major improvements of the model.

The equations for chloride dynamics and the parameters used are summarized in Appendix B; their integration into the NMM equations is illustrated by Figure 3.

### 2.6. Personalization of the model

#### 2.6.1. Dynamical landscape

The set of NMM parameters selected corresponds to a parameter space where different epileptic transitions can be achieved by varying the synaptic gains of the connection between populations of inhibitory interneurons and pyramidal populations (see Figure 4). While most parameters in the model inherit from previous work [1], some are personalized to reproduce the main features of the patient SEEG signal in the contact recording the most epileptogenic activity.

**Figure 4:**
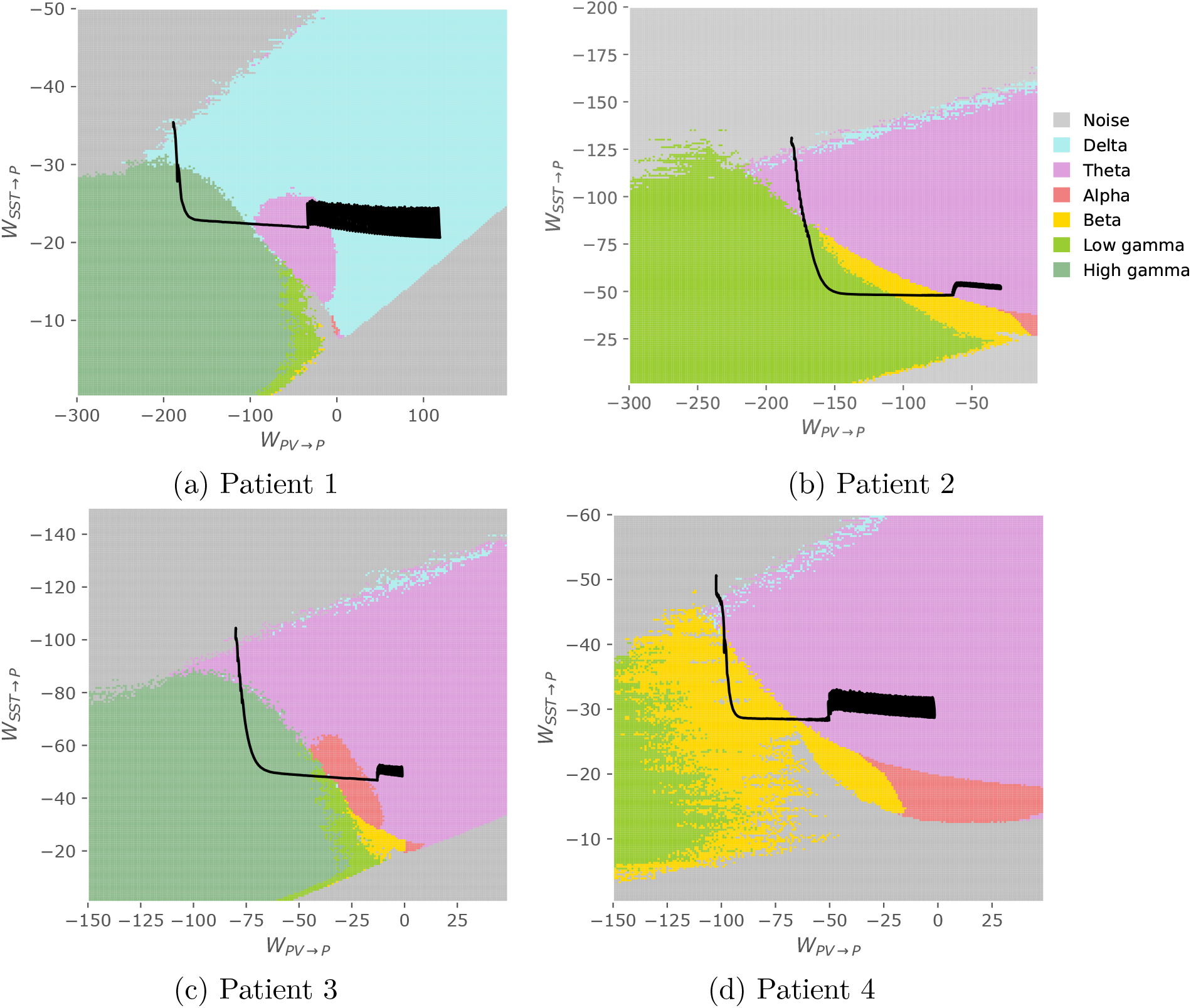
Dynamical landscape of the model corresponding to different inhibitory synaptic weights. Different types of epileptic activity can be found for different values of the pair *W*_SST→P_, *W*_PV→P_: background activity (grey), spiking in the delta (light blue) or theta (light pink) range, rhythmic oscillatory activity in the alpha (red) or beta (yellow) band, low voltage fast activity in the low-gamma (light green) or high-gamma (dark green) band. The solid black line illustrates the path taken in the landscape, determined by chloride dynamics.

In general, the parameter space must include the following regions: background activity, fast oscillatory activity (if the patient’s SEEG recordings display fast onset activity) and slow rhythmic activity. The main frequency of each of those regions should correspond approximately to the ones extracted from the patient’s data.

The adjustment of parameters to achieve the desired parameter space is performed starting from the set of parameters presented in [1] and modifying as few parameters as possible‡. Some parameters are prioritized from physiological considerations, including the synaptic gain of excitatory connections (which reflects the excitability of the epileptic tissue) and the connectivity strength of synapses (some connections might be strengthened or weakened due to the frequent epileptic activity in the region).

The parameters that define population average PSP kinetics are instead expected to remain within realistic ranges, which we have defined based on previous modeling studies [1, 39, 30]. In particular, the work of [39] provides literature-based intervals for the time constants of glutamatergic and GABAergic PSPs. The ranges of PSP rates used in our study are 100 to 220 s^−1^ for *P* and *E* pre-synaptic populations, 125 to 500 s^−1^ for *PV* pre-synaptic populations, and 15 to 50 s^−1^ for *SST* pre-synaptic populations.

Figure 4 describes the resulting parameter space of all patients, and Table 2 provides the set of parameters chosen for the personalized model of each patient. Different types of epileptic activity (background activity, rhythmic oscillatory activity, low voltage fast activity) can be found for different values of the pair *W*_SST→*P*_, *W*_PV→*P*_ (see Figure 4). The parameters defining the sigmoid for the translation of membrane potential perturbation into firing rate (*φ*_0_, *r, v*_0_) were fixed for all neuronal types to the values presented in [30]: *φ*_0_ = 2.5 Hz, *r* = 0.56 mV^−1^, *v*_0_ = 6 mV. The external input to the pyramidal population was for all patients Gaussian noise with a mean of 90 s^−1^ and a standard deviation of 30 s^−1^.

**Table 2:** NMM synaptic parameters chosen for all patients. For synaptic gains (*W*) and PSP rates (*τ*), subscripts indicate the type of pre-synaptic population (*exc* corresponds to both P and E populations). Some parameters are taken from [1], while others have been adjusted to obtain the patient-specific parameter space that includes the transitions phases required to fit SEEG data. Parameters that have been modified with respect to the values used in [1] are marked with an asterisk. In this case, the synaptic gain of excitatory connections has been increased (reflecting an increase of excitability), along with some synaptic connectivity values and the PSP rates of some synapses.

| Parameter | Patient 1 | Patient 2 | Patient 3 | Patient 4 |
| --- | --- | --- | --- | --- |
| $W_{exc}$ [mV] | 20* | 15* | 20* | 7* |
| $W_{SST}^a$ [mV] | -22 | -22 | -22 | -22 |
| $W_{PV}^a$ [mV] | -10 | -10 | -10 | -10 |
| $1/\tau_{exc}$ [s<sup>-1</sup>] | 180* | 100 | 180* | 100 |
| $1/\tau_{SST}$ [s<sup>-1</sup>] | 20* | 50 | 50 | 50 |
| $1/\tau_{PV}$ [s<sup>-1</sup>] | 500 | 500 | 500 | 500 |
| $C_{E \rightarrow P}$ | 108 | 108 | 108 | 108 |
| $C_{SST \rightarrow P}$ | 33.75 | 33.75 | 33.75 | 33.75 |
| $C_{PV \rightarrow P}$ | 108 | 108 | 108 | 108 |
| $C_{Ext \rightarrow P}$ | 1 | 1 | 1 | 1 |
| $C_{P \rightarrow E}$ | 135 | 135 | 135 | 135 |
| $C_{P \rightarrow SST}$ | 33.75 | 33.75 | 33.75 | 33.75 |
| $C_{P \rightarrow PV}$ | 150* | 40.5 | 150* | 40.5 |
| $C_{SST \rightarrow PV}$ | 3* | 3* | 3* | 3* |
| $C_{PV \rightarrow PV}$ | 800* | 300* | 450* | 300* |
| $v_0$ [mV] | 6 | 6 | 6 | 6 |
| $\varphi_0$ [s<sup>-1</sup>] | 2.5 | 2.5 | 2.5 | 2.5 |
| $r$ [mV<sup>-1</sup>] | 0.56 | 0.56 | 0.56 | 0.56 |
| $p_m$ [s<sup>-1</sup>] | 90 | 90 | 90 | 90 |
| $p_{std}$ [s<sup>-1</sup>] | 30* | 30* | 30* | 30* |

\* Values modified with respect to [1]
<sup>a</sup> $W_{SST \rightarrow P}$ and $W_{PV \rightarrow PV}$ are [Cl<sup>-</sup>]-dependent

#### 2.6.2. Chloride accumulation parameters

Once the parameters defining the dynamical landscape are selected for each subject, we fit the parameters describing the chloride accumulation dynamics. These parameters determine how the pair of parameters (*W*_SST→*P*_, *W*_PV→*P*_) moves in the dynamical landscape as chloride accumulation evolves, driving the system through different types of epileptic activity. The goal of the fit is to approximate the transition from interictal to seizure found in the SEEG recordings of each patient. The evolution of *W*_SST→P_ and *W*_PV→P_ in the personalized dynamical landscape of all patients is shown in Figure 4.

To reduce the dimensionality of the problem, we have fixed the values of *α*_KCC2_ and *α*_*φ*_ to 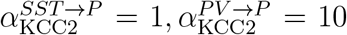 and 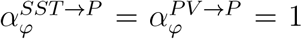. These parameter values lead to [Cl^−^] equilibrium values in inter-ictal state of around 11 mM in the dendrites and 8.5 mM in the soma. These are above typical healthy values, and are consistent with measured [Cl^−^] in KCC2-impaired neurons of animal models [40]. As mentioned above, we hypothesize that the concentration of chloride is larger in the dendrites due to the lower volume of the compartment and limited diffusion capacity.

We are left with four key parameters that regulate the chloride accumulation in the synapse locations (see Table 3). Our first step was to personalize the parameters *w*_0_ and *W*_*h*_, used to compute the dynamic synaptic gains (*W*_SST→*P*_, *W*_PV→*P*_) from GABAergic currents. The values of *α*_KCC2_ and *α*_*φ*_ (in the two synaptic locations) determine the equilibrium point of chloride accumulation for a given firing rate. Those parameters being fixed, each activity type in the dynamical landscape will lead to an equilibrium value of *E*_GABA_. Thus, choosing *w*_0_ and *W*_*h*_ to achieve the desired transitions amounts to solving a simple system of two equations (Equations 15 and 16). A description of the process can be found in Figure S4.

**Table 3:**
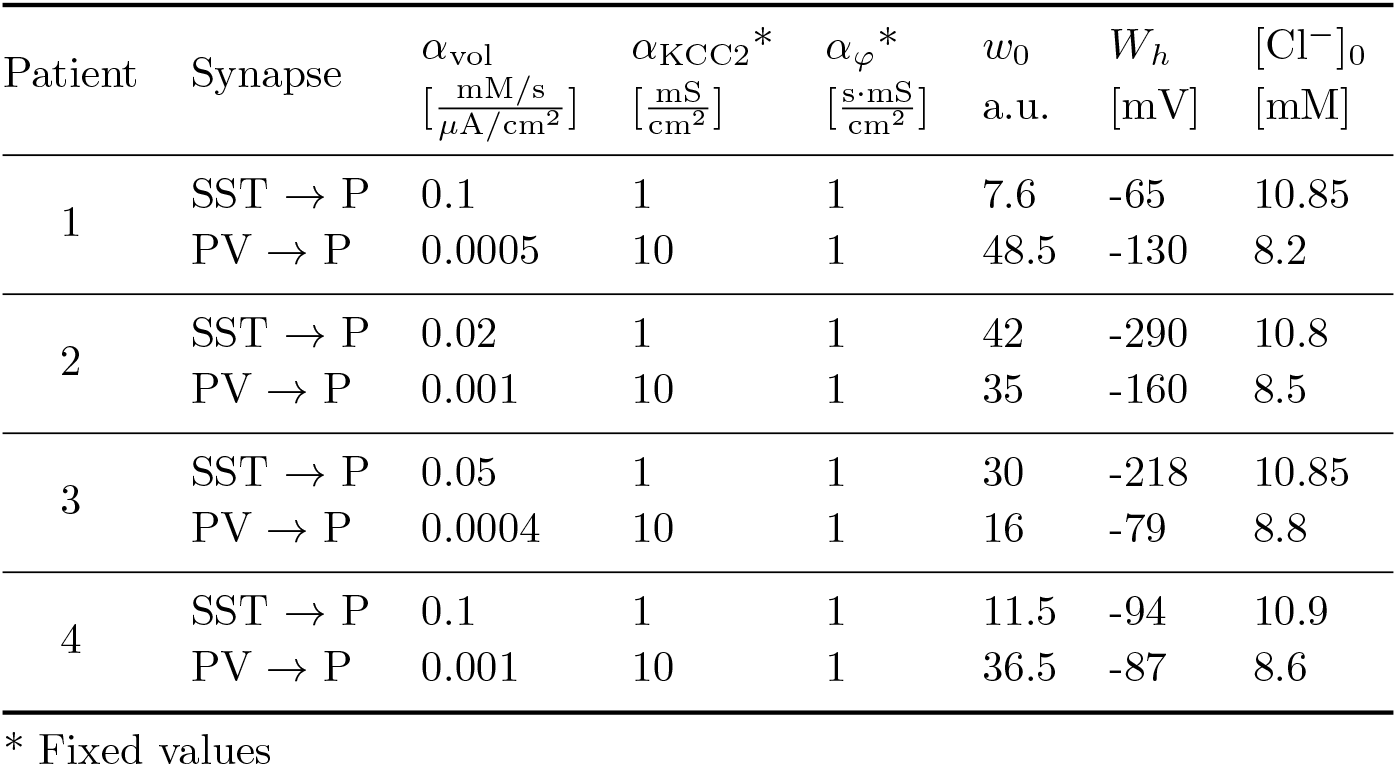
Chloride dynamics parameters. The parameters regulating the chloride accumulation in the pyramidal population are adjusted so that the evolution of synaptic gains in the parameter space drives the system to the different transition phases observed in the patient’s SEEG data.

Then, we personalized the parameter *α*_vol_, which regulates the rate of chloride accumulation in the different compartments. Following the assumption that chloride accumulates faster in apical dendrites than in the soma, we require 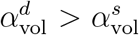. These parameters are used to adjust the duration of the different phases and ensure that transitions match those observed in the patient data. Finally, we have chosen [Cl^−^]_0_ so that it corresponds to the equilibrium value of [Cl^−^] in the inter-ictal state.

As an illustrative example, Table 3 summarizes the values of chloride dynamics parameters selected for all patients.

Regarding the rest of variables determining the dynamics of chloride accumulation, the concentration of 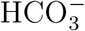 inside and outside the cell was fixed at constant values, as well as the concentration of chloride outside the cell: 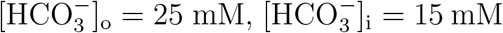, [Cl^−^]_o_ = 150 mM. We also assumed the reversal potential of potassium and the average membrane potential of the cell to be constant (needed for the calculation of KCC2-driven chloride transport and for the calculation of GABAergic currents respectively): 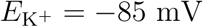, *V*_m_ = −65 mV.

As mentioned in the previous section, the parameter *α*_KCC2_ represents the degree of pathology in the simulated node. Given that an increase of this parameter corresponds to improved KCC2 transport, we tested and confirmed that increasing *α*_KCC2_ above certain values forces the system to remain in non-pathological background activity (see Appendix E). Similarly, decreasing the average value of the external input leads to a decrease of the excitation/inhibition ratio in the NMM. Since the transition to epileptic seizure is driven by stochastic fluctuations of the external input to the pyramidal population, we verified that by setting the external input to sufficiently low values, the transition to ictal state is avoided (more details are provided in Figure E1).

### 2.7. Physical model for generation of SEEG data from NMM results

In order to compare experimental data with our modeling results, it is necessary to connect the NMM formalism with a physical modeling framework. In the present study, we built a model to estimate the bipolar voltage recorded by a pair of consecutive contacts in an SEEG electrode from the laminar NMM source activity. A detailed description of the model can be found in Appendix C.

In brief, we considered that the synapses targeting the pyramidal population are the main contributors to the measured voltage, given the anatomical characteristics and organization of pyramidal cells [41, 42]. In the model, the pyramidal population spans from layer V to layer I, and we have assumed that the synapses targeting this population are located in either apical (layer I) or basal (layer V) locations (see Figure 2, and Section 2.4). For each of these two locations, we built a simplified current source distribution model with discrete sources to approximately reproduce the current distribution from compartmental models. These sources were embedded in a three layer axisymmetric model of a cortical patch with a realistic representation of the SEEG contacts. Electrical conductivity values to represent the white matter, gray matter and cerebro-spinal fluid were assigned to these layers from the literature [43], and the electric potential distribution was solved using the finite elements method (FEM).

The results from the FEM model described above were used to calculate the relative contribution of synaptic currents in each location to electrical potential as a function of the SEEG contact depth with respect to the cortical surface. With this information, the voltage difference over time measured by a pair of SEEG contacts can be approximated as the weighted sum of the aggregated synaptic current in each NMM synapse *I*_*s*_:

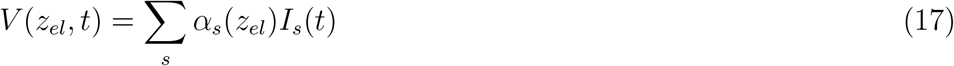

where *α*_*s*_(*z*_*el*_) has units of V/A and is the relative contribution of each synaptic current to the total voltage (from the FEM model). The value of *α*_*s*_(*z*_*el*_) depends on the SEEG contact depth *z*_*el*_ as well as the location of the synapse. Note that in this study only two possible locations were considered (apical and basal). In line with our earlier simplifying assumptions equating synaptic current and potential time constants, we take the post synaptic current *I*_*s*_(*t*) as proportional to the synapse membrane potential perturbation *u*_*s*_(*t*) — a direct output of the NMM. The proportionality factor depends on the post-synaptic neuron morphology and membrane conductivity, and is represented by a factor *η*_*n*_ (A/V). Finally, to match the features of real SEEG data, simulated SEEG signals are high-pass filtered above 0.5 Hz.

## 3. Results

### 3.1. A realistic model of interictal to ictal transition

The model generates autonomous realistic transitions to ictal activity. As described in the *Methods* section, this is achieved by adding dynamical equations to the NMM to account for Cl^−^ transport and its subsequent impact on inhibitory post-synaptic potential (PSP) amplitude. In the model, realistic seizures are generated when homeostatic mechanisms of pyramidal cells are dysfunctional (KCC2 transporter dysregulation) and unable to cope with the influx of Cl^−^ from pathological excitation of GABAergic inhibitory cells such as SST or PV interneurons.

The cascade of events leading to the transition from interictal to ictal state reproduced by the model is illustrated in Figure 5 and can be summarized as follows:

**Figure 5:**
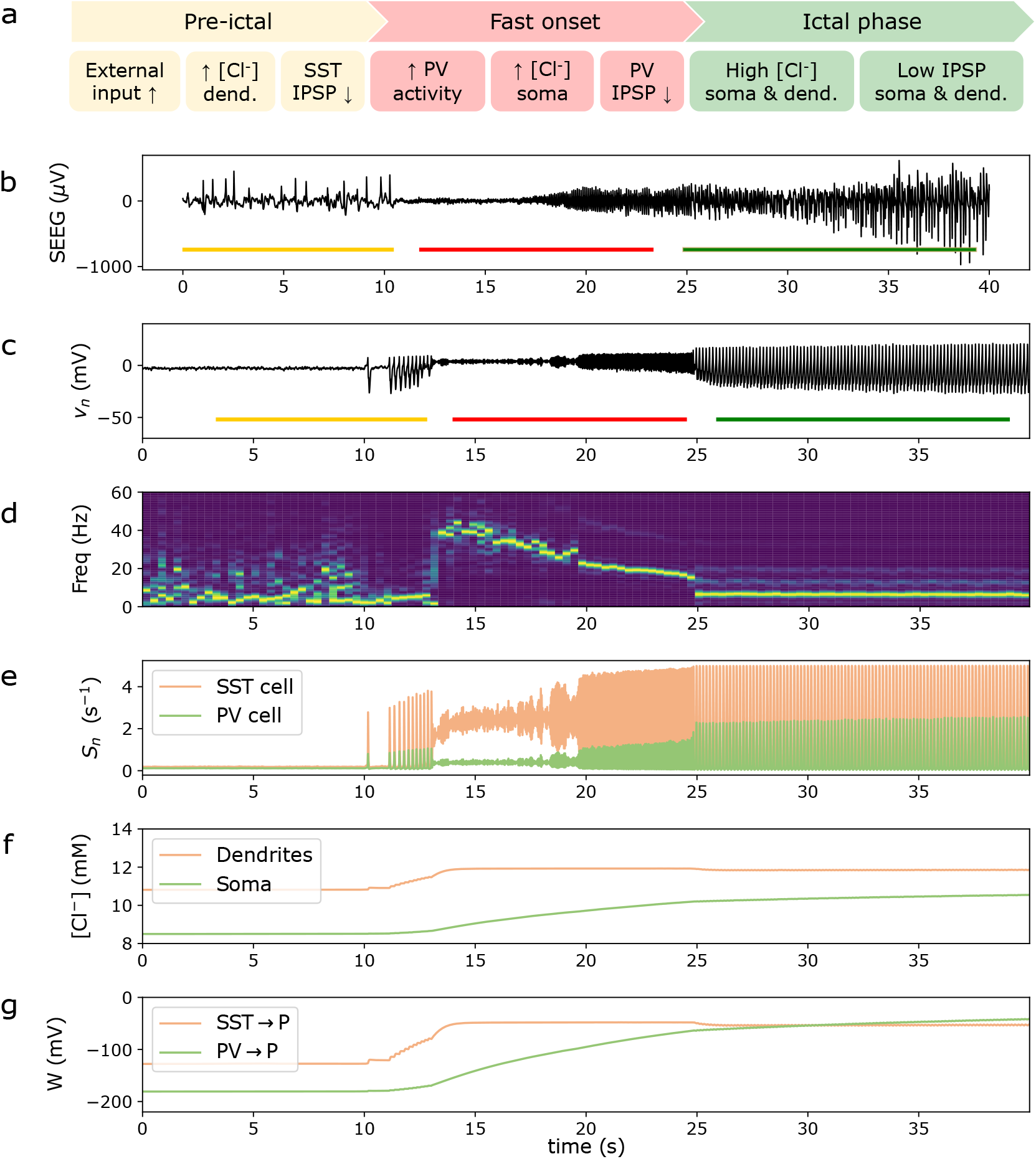
A realistic spontaneous seizure generated by the the model. (a) Summary of the cascade of events leading to the transition from interictal to ictal state. (b) Real SEEG signal recorded during a seizure of Patient 2. (c) Time series of the pyramidal cell population average membrane potential, and (d) weighted time-frequency representation of the signal. Different transition phases are clearly observed: interictal activity, pre-ictal spiking, low voltage fast activity and rhythmic ictal activity. (e) Firing rate of the GABAergic inhibitory interneuron populations (SST and PV). (f) Evolution of Cl^−^ accumulation in the two pyramidal cell compartments represented in the model (dendrites and soma). The pathological accumulation of Cl^−^ in the cells drives the system through different phases of the seizure. (g) Evolution of the average synaptic gain (IPSP amplitude) of the synapses from inhibitory interneuron populations (PV and SST) to pyramidal cells. The simulation corresponds to the model with parameters adjusted using Patient 2 data.

i. Random fluctuations in the external excitatory input to the main pyramidal cell population lead to an increase in excitation.
ii. This increase in excitation produces an increase of activity of SST interneurons, which leads to Cl^−^ overload in the dendritic compartments of the pyramidal cell population (P). Due to the dysfunction of the Cl^−^ transporter system, Cl^−^ cannot be extruded sufficiently fast and accumulates inside the dendrites (Figure 5(f), orange curve).
iii. The accumulation of Cl^−^ in dendritic compartments leads to a decrease of the inhibitory PSP (IPSP) amplitude generated by GABA release from SST cells. In the model, this translates into a decrease of the magnitude of the synaptic gain of GABAergic synapses from SST to P cells, *W*_SST→P_ (Figure 5(g), orange curve), and even its reversal (*W*_SST→P_ and *W*_PV→P_ are *signed* gains in the model, e.g., they are negative when inhibitory).
iv. The consequent reduction of inhibition leads to a drift of system dynamics, which produces epileptic spikes (Figure 5(c), period marked in yellow) followed by low-voltage fast activity (Figure 5(c), period marked in red), a typical marker of the onset of seizure during the tonic phase [24, 44, 45]. The fast activity is mainly generated by the fast loop circuitry associated with the PV population (see Figure 2), constituted by the autaptic connections among parvalbumine-positive cells (PV) [46] and the connection between PV and pyramidal neurons.
v. The increased activity of PV cells leads to Cl^−^ overload in the somatic compartment of the pyramidal population, which accelerates Cl^−^ accumulation in the soma due to the limited extrusion capacity of KCC2 transporters (Figure 5(f), green curve).
vi. An increase of somatic [Cl^−^] in the pyramidal cell leads to a decrease of the IPSPs generated by the GABAergic neurotransmitters released by PV cells, which translates in the model into a weakening of the synaptic gain and even its reversal (Figure 5(g), green curve).
vii. The reduction of fast inhibition leads to a new drift of system dynamics, producing the rhythmic ictal activity usually observed during the clonic phase (Figure 5(c), period marked in dark green).

Figure 5 provides a sample model time series of the average membrane potential of the pyramidal cell population along with the corresponding time-frequency representation during a spontaneous seizure. The signal reproduces the temporal dynamics typically observed in SEEG seizure recordings: pre-ictal spikes, low-voltage fast activity and transition to rhythmic slow activity in the ictal phase. In particular, the simulated seizure is generated by the model with the parameters adapted to Patient 2. A key feature of the model is also to simulate the firing rate of GABAergic inhibitory interneurons (PV and SST), the evolution of Cl^−^ accumulation, as well as the evolution of the synaptic gains of GABAergic synapses. Once a model is personalized, access to these dynamical variables can provide valuable clinical insights.

### 3.2. Patient-specific personalized models

Model parameters can be personalized to fit the main features of a specific SEEG recording. In particular, we focused on reproducing the frequency of the different phases of the recording during the interictal to ictal transition. The fitting takes place in a two-step process: first, the NMM parameters are adjusted to obtain a *dynamical landscape* that includes the dominant SEEG frequencies found during the different seizure phases. The choice of focusing on the representative frequency of each transition phase for model fitting, instead of other SEEG features such as spike morphology, is driven by considerations about SEEG physics that are discussed in the next section. The landscape summarizes the dynamics associated with model parameters. Here we focus on a two dimensional slice corresponding to the (dynamical) synaptic gains of inhibitory interneurons, *W*_SST→P_ and *W*_PV→P_, with all the other parameters fixed (see Figure 4).

In a second step, the parameters regulating the dynamics of chloride accumulation are adjusted. These parameters determine the particular path the system takes in the dynamical landscape of inhibitory synaptic gains, driving the system through the different phases of the seizure. Further details about the chloride dynamics parameters and the model fitting process are given in the *Methods* section. As illustrative examples, Figure 4 displays the path traversed by the gain variables in the landscape map corresponding to the personalized NMM model for all patients. In all cases we used a slight variation of the parameters (other than *W*_SST→P_ and *W*_PV→P_) found in [1]. The set of NMM parameters used for all patients is provided in Table 2.

Crucially, the pipeline presented in this study allows for personalization of a wide variety of features found in SEEG recordings during seizure: high-frequency fast onset (Patients 1 and 3), low-frequency fast onset (Patient 2) or no fast onset (Patient 4); rhythmic ictal activity in the delta-range (Patient 1) or theta-range (Patients 2, 3 and 4); and presence of pre-ictal spikes (Patients 2 and 3) or not (Patients 1 and 4).

Figure 6 illustrates how these patient-specific models, obtained with modifications of a few parameters (see Table 2), match the most representative frequencies of each seizure transition phase found in the SEEG data. For a given patient, these representative frequencies display a certain variability across seizures, which is more evident in the rhythmic ictal phase (Figure 6, right panel). It is worth mentioning that this variability could also be partially reproduced by the personalized models, notably by altering the standard deviation of the external input noise that drives the model into seizure. However, given the limited seizure data available (maximum four seizures per patient in our dataset), we have not focused on reproducing this aspect of the data.

**Figure 6:**
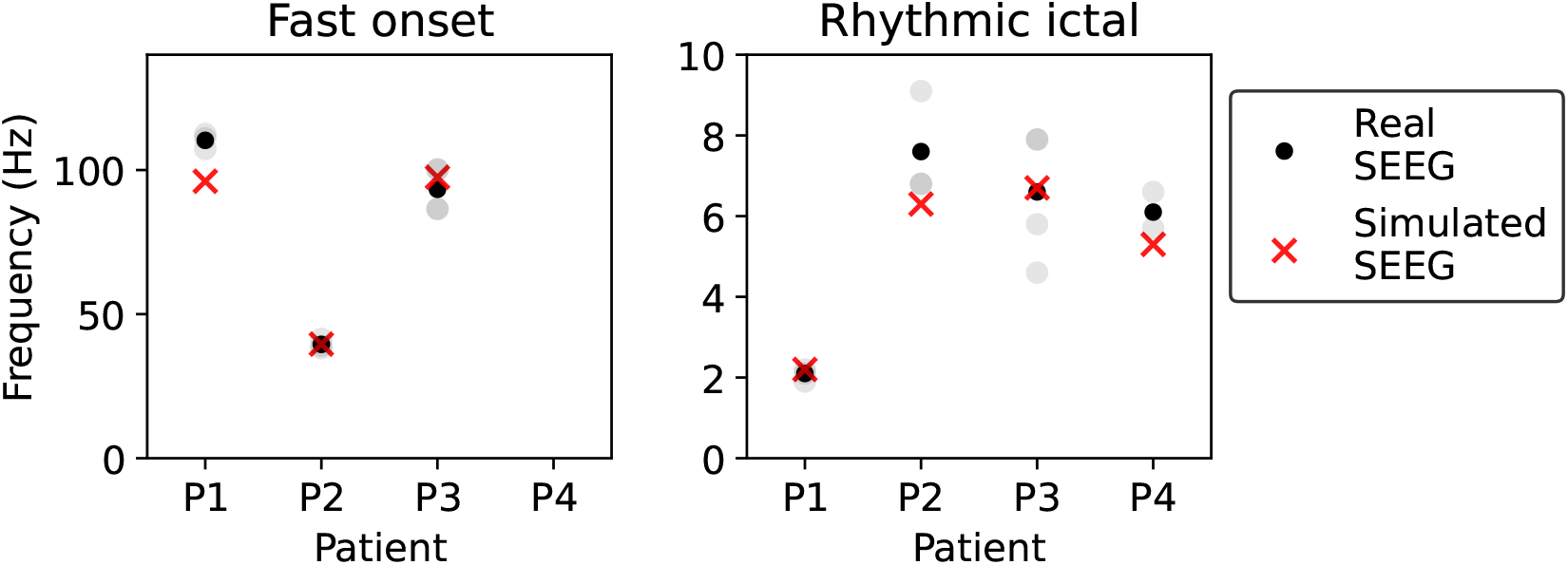
Real and simulated main frequencies of each seizure phase for each of the four patients. For each patient and seizure, we have selected the periods of low voltage fast onset and rhythmic ictal activity, and computed the representative frequency of each of these phases (see *Methods* for more details) with the aim of reproducing them in the personalized model. Three SEEG seizure recordings were available for Patients 1 and 2, four for Patient 3 and two for Patient 4. The representative frequency values for each patient are computed as the average over the total number seizure recordings. Grey dots represent the main frequency in the fast onset or rhythmic ictal phase for a single seizure and black dots represent the average value over seizures. The variability between seizures is lower in some patients (e.g., Patient 1) and seems more prominent in the rhythmic ictal phase. Red crosses indicate the representative frequency for each patient and seizure phase in the simulated data. The model represents accurately the relevant frequencies of the fast onset and rhythmic ictal phases observed in the patient data.

The time periods selected as representative of the low voltage fast activity (LVFA) phase and the rhythmic phase in each patient’s seizures are shown in Figures S3 (real data) and S4 (simulated data). Table D1 provides a detailed comparison of different SEEG features for simulated and real data of all patients.

### 3.3. Simulating SEEG contact activity in the epileptogenic zone

As described in the *Methods* section, the laminar NMM is embedded into a physical model in order to produce realistic SEEG data. The physical model places the cell subpopulations and synapses (current sources) in neocortical layers using a local representation of conductive media. The laminar architecture allows for the representation of the activity generated in each cortical layer of the epileptogenic zone. The physical forward model translates synaptic current source activity in each layer into a realistic SEEG signal on the desired SEEG electrode contact by solving Poisson’s equation.

The morphology of SEEG signals in our model is highly sensitive to the precise position of the electrode contact, specially in terms of depth inside the grey matter. For the present study, we could not determine the electrode position inside the grey matter with enough resolution from the computed tomography (CT) scan images. To overcome this limitation, we have simulated SEEG signals for different possible locations of the most epileptogenic contact. An example of simulated SEEG signals for different electrode depths for a personalized model (Patient 3) can be found in Figure 7.

**Figure 7:**
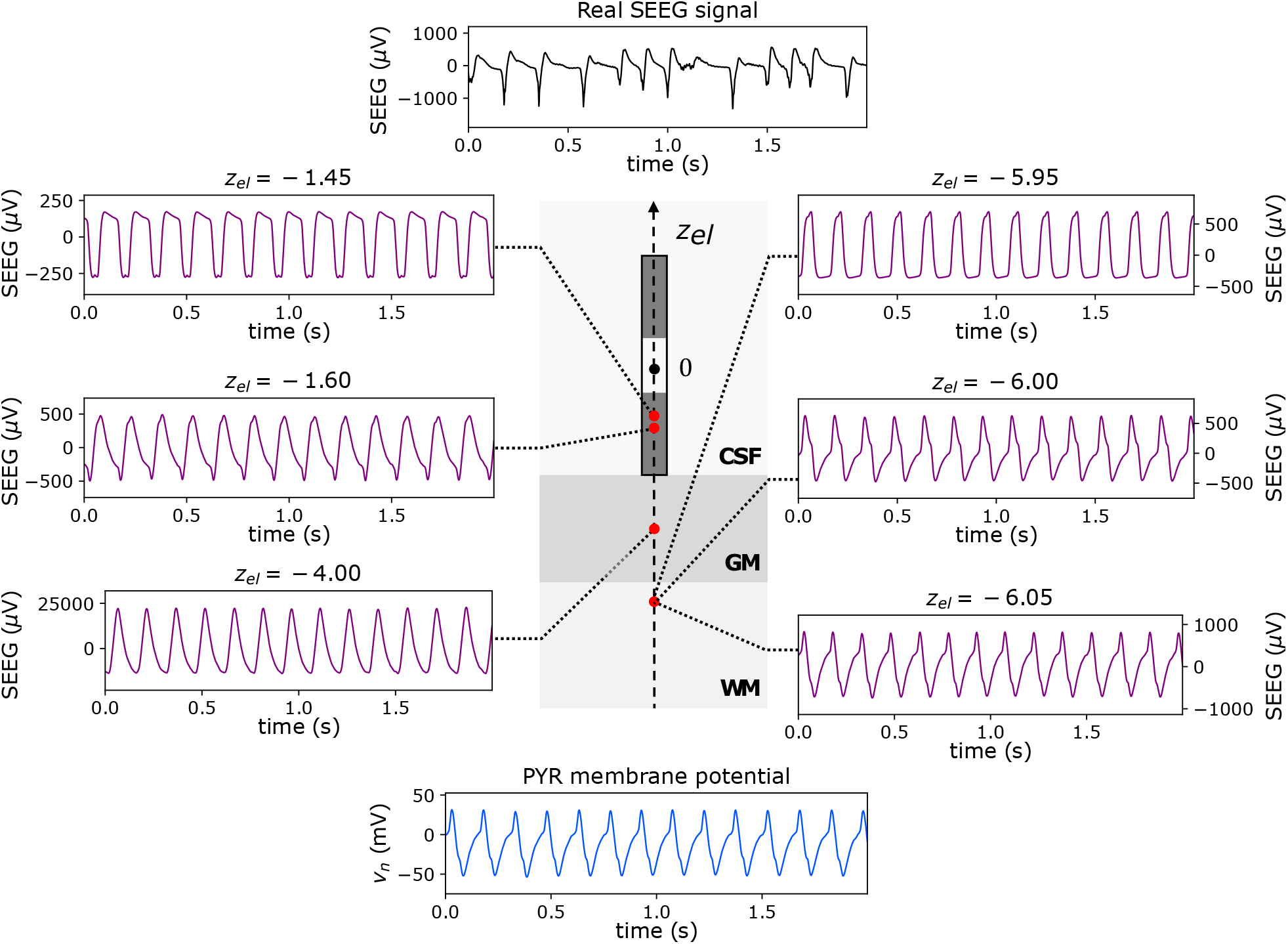
Different electrode depths lead to different SEEG morphology, and some electrode positions approximate the real SEEG signal morphology. The main frequency is preserved for different electrode positions, and for the membrane potential perturbation of the pyramidal population. *Central figure*: Schematic view of the physics axisymmetric model used to generate realistic SEEG signals (see Figure C2 for more details). Only part of the SEEG lead has been modeled, including two contacts for bipolar measurements (represented in dark grey). We obtain different signals for different depths *z*_*el*_ of the electrode (see other panels); this schema illustrates the case *z*_*el*_ = 0 mm, where the bottom electrode contact is exactly above the grey matter, both electrode contacts being in the CSF region. *Left panels*: Morphology and amplitude of the signal depend strongly on the electrode depth (notice the changes of scale in the bottom panel). Some electrode depths lead to signal morphologies that resemble the ones observed in the real SEEG data (shown in the top panel), in this case for *z*_*el*_ = −1.6 mm (middle left panel) a good approximation is found. *Right panels*: very small variations of the electrode depth in the physical model lead to substantial changes in signal morphology. *Bottom panel* : The membrane potential perturbation of the main pyramidal population can approximate the SEEG signal morphology only for very specific depths (in this case, for *z* ≈ −6.05 mm, see bottom right panel), but the main frequency is captured. All plots correspond to the ictal period of Patient 3. *z*_*el*_ units are mm.

As predicted, the signals display a wide variety of spike morphologies as a function of contact depth, some of which approximate well the real SEEG recordings. The signal morphology changes substantially with small variations of the electrode depth (as small as 50 *µ*m, see right panels), which confirms that the precision of the CT scan used to retrieve the electrode position inside the brain (of the order of 0.5 mm) was not sufficient for the purpose of morphology modeling. Moreover, given the variability of signal morphologies for different electrode depths, the membrane potential perturbation of the main pyramidal population is not a good surrogate of the SEEG signal in terms of morphology. In fact, it is only representative of signal morphology for very specific depths (in this case, for *z* ≈ −6.05 mm, see bottom right panel).

In contrast, the representative frequency of each transition phase is preserved across all contact positions, and is also well reproduced by the membrane potential perturbation of the pyramidal population. These results justify our two-step modeling strategy: first, the NMM parameters were adjusted so that the spectral features of the NMM output— represented by the membrane potential perturbation of the pyramidal population— matched the ones found in real SEEG recordings (see Section 3.2). In a second step, we added a physical layer to the laminar NMM for the simulation of realistic SEEG signals from the personalized model. In this case, we heuristically fitted the electrode depth so that the morphology of simulated SEEG signals could match the one of real data.

Figure 8 displays the SEEG electrode signal recorded during one seizure for all patients, compared with the respective simulated signals obtained from the laminar NMM embedded in the physical head model. Although our main focus was to match the main frequencies in each phase, we present some simulated SEEG signals corresponding to an electrode location that provides a good fit in terms of signal morphology. Figures D1 and D2 provide further examples where the morphology of the real signal can be approximated by the simulated SEEG signal at a given electrode depth.

**Figure 8:**
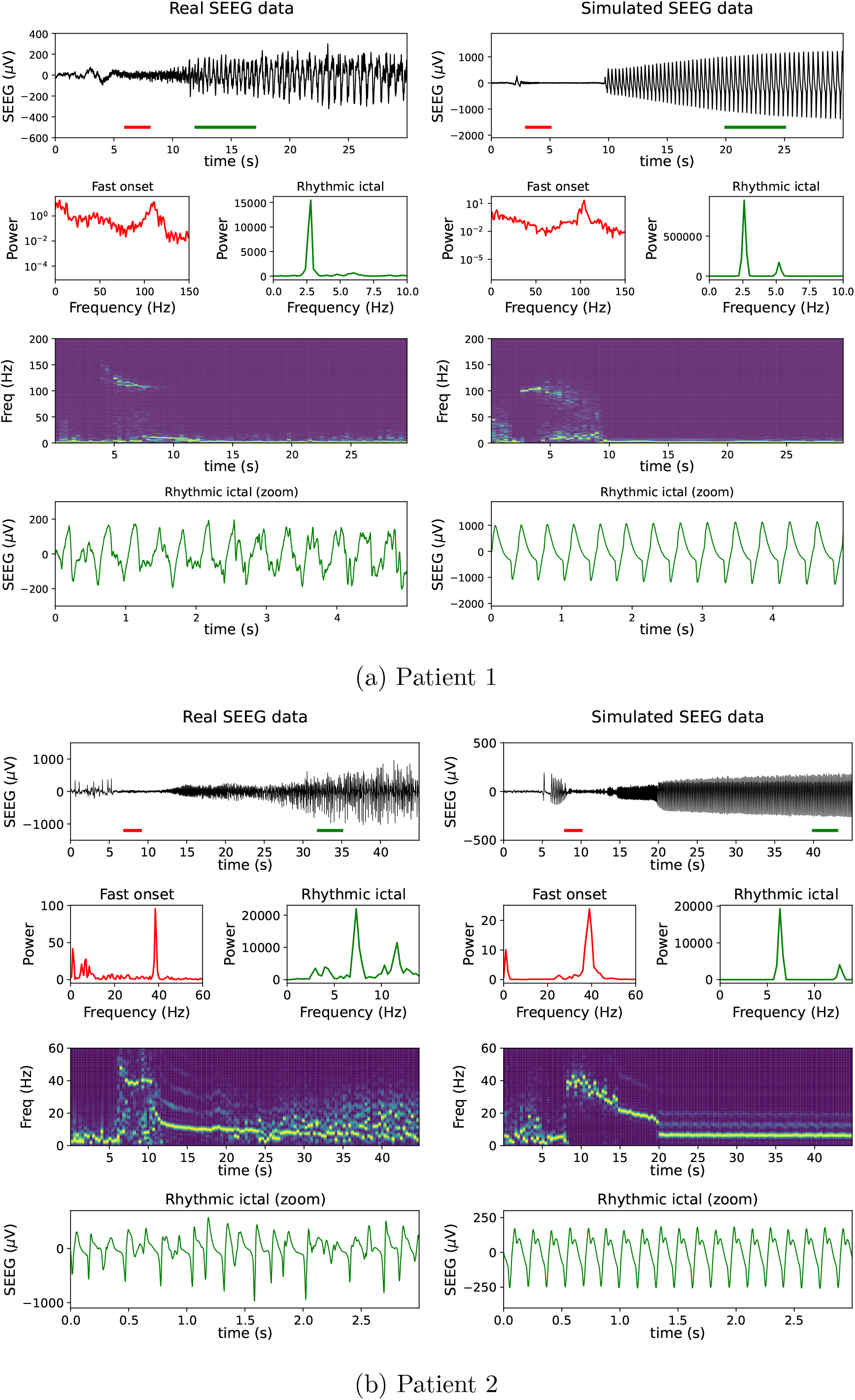
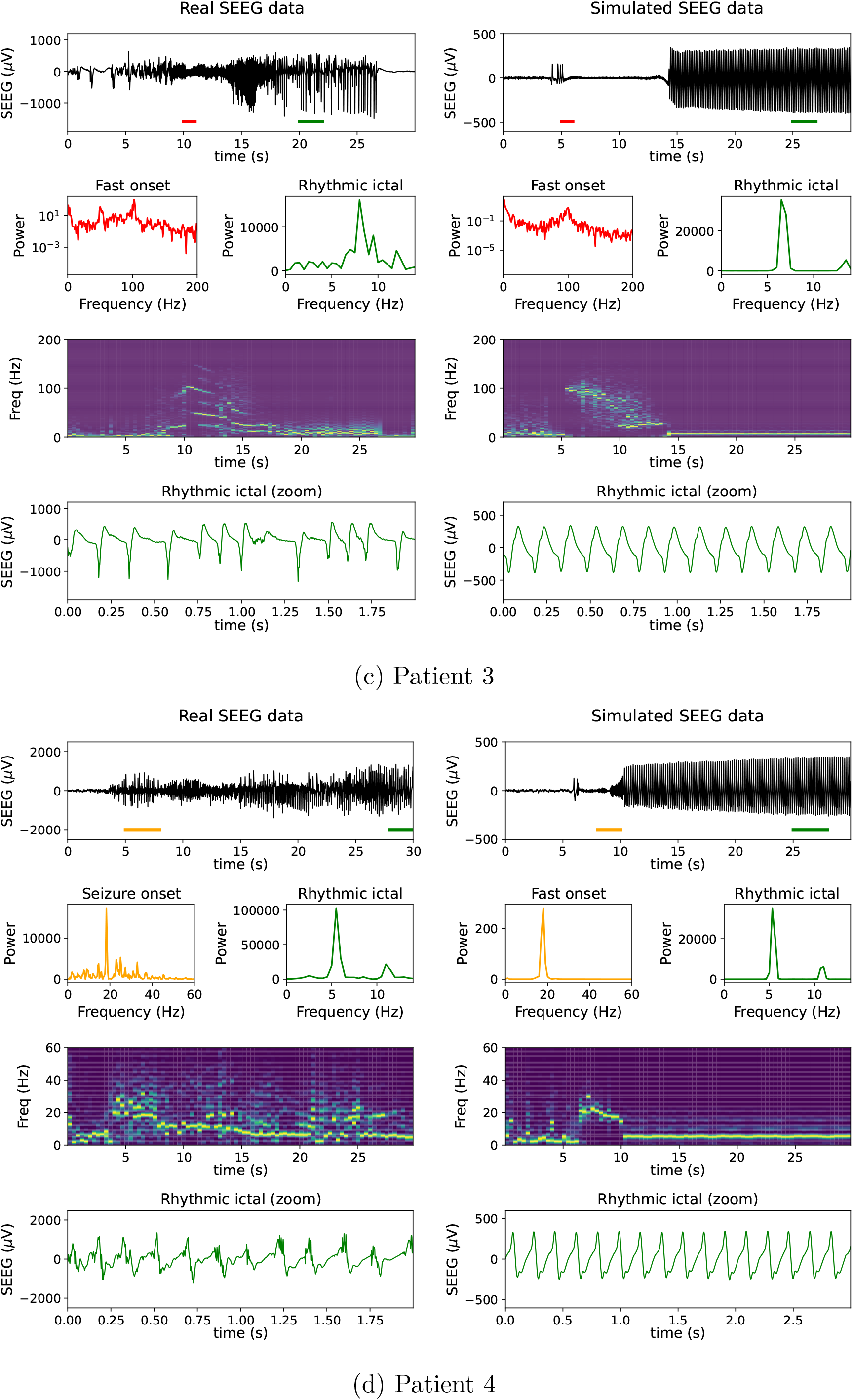
Generation of realistic SEEG signals with personalized models. Panels on the left display SEEG recordings in the most epileptogenic contact, and panels on the right display the simulated SEEG from the personalized model. The red and green traces in the SEEG time series indicate the periods selected for power spectral density (PSD) analysis of the LVFA and rhythmic phases respectively, shown below. For Patient 4 no LFVA phase was detected, but a frequency higher than in ictal phase (beta range) was observed at seizure onset. We have also computed the PSD for this phase, marked with an orange trace. For Patients 1 and 3, a logarithmic scale has been used in the fast onset PSD representation to capture the 100 Hz peak frequency at seizure onset. Below the PSD plots, a weighted time-frequency representation of the signal is shown, which captures the high-frequency traces at seizure onset and the transition to rhythmic activity (see *Methods*). The bottom panels provide a zoomed view of the signal during the ictal phase. The simulated SEEG signals have been produced by embedding the patient-specific laminar NMM in a physical cortical model. Given that different electrode depths lead to very different signal morphology (see Figure 7), our main focus has been to fit the main frequency in each phase of the transition from inter-ictal to seizure. The signals displayed in the bottom panels correspond to the particular locations of the SEEG electrode in the physical model that provide a good representation of the signal morphological features.

### 3.4. Stimulation of epileptogenic nodes: proof of concept

To better understand the effect of transcranial electric stimulation (tES) for the treatment of epilepsy, we have simulated the effect of electric fields in the personalized model of the epileptogenic node of all the patients in the study. As explained in the *Methods*, the NMM synapse equations include the perturbation induced by an electric field in the average membrane potential of a neuronal population (Equation 3). The term *λ*_*n*_*·E*(*t*) represents the membrane perturbation induced by an external electric field. The coupling term *λ*_*n*_ used — which represents the membrane perturbation induced by a unitary external electric field — has been set for convenience to *λ*_*n*_ = 0.1 mm, a value found in the experimental [47] and modeling literature [48].

Figure 9(a) displays the simulated SEEG signal for different values of the electric field applied to the personalized model of Patient 2. It can be seen that for an inhibitory electric field of amplitude 0.2 mV/mm the seizure completely disappears, while for 0.1 mV/mm its appearance is delayed. The inhibitory electric field of 0.2 mV/mm represents the average electric field (normal component) stimulation on a small cortical patch containing the epileptogenic node with cathodal transcranial direct current stimulation (tDCS) of amplitude 1 mA [31].

**Figure 9:**
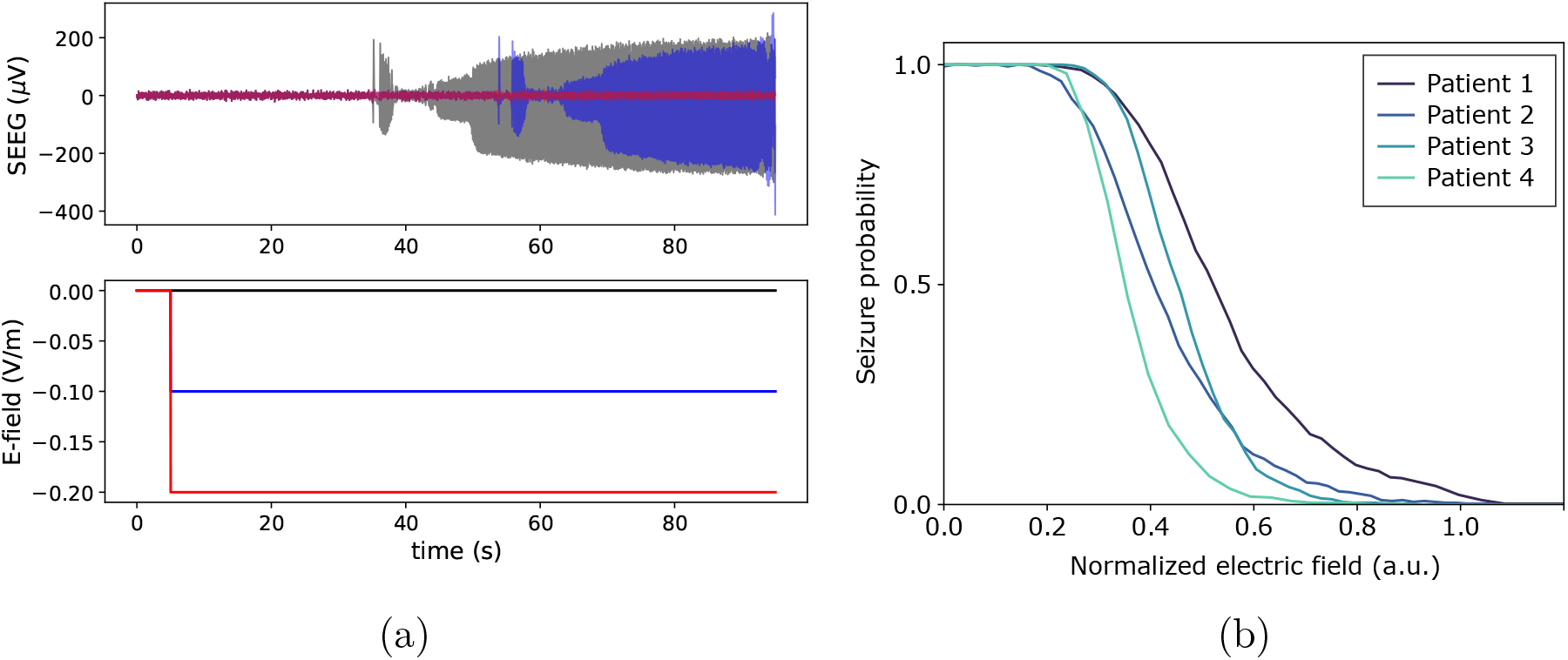
Simulation of tDCS effects in epileptogenic nodes. (a) Example with the personalized model of Patient 2. The top panel displays the simulated SEEG signal (*z*_*el*_ = −1.55 mm) for three different cathodal stimulation scenarios, shown in the bottom panels: no stimulation (black), 0.1 V/m (blue), and 0.2 V/m (red). The latter is sufficient to avoid the seizure event. (b) Reduction of seizure probability in the personalized models as a function of the electric field applied. The electric field perturbation has been normalized to homogenize the results across patients (see main text). The seizure probability is computed as the probability to detect a seizure in a time window of 100 seconds over 500 simulations. Each personalized model responds differently to the electrical stimulation, and some models (e.g., for Patient 4) require a lower intensity of the electric field to achieve complete seizure suppression.

We have also studied the reduction of seizure probability for all the personalized models as a function of the electric field applied. To homogenize the results across patients, the effect of the electric field is normalized by the standard deviation of the pyramidal population membrane potential perturbation in interictal state. Specifically, 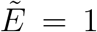 a.u. corresponds to an electric field that changes the pyramidal population the equivalent of 1 std of the baseline membrane potential. The seizure probability represents the probability of finding a seizure within a 100 seconds time window for 500 different realizations of the external input (Gaussian white noise)—we recall that the model is driven into seizure by the stochastic fluctuations of the external input to the pyramidal population. The standard deviation of the external input was increased to 40 s^−1^ in all the models to adjust the seizure probability without stimulation to 1. We selected a 100 seconds window so that the model would display seizures consistently in the no-stimulation case, while keeping the computational cost low.

Figure 9(b) shows the probability curves as a function of the normalized electric field amplitude for the four patient models. It can be seen that there is a minimum amplitude of the electric field needed for any seizure reduction, which is evidenced by the initial flat segment on the probability curve. For instance, for Patient 4, the probability starts decreasing around 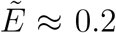 (a.u.). After this initial part, the seizure probability reduction is gradual, and follows a sigmoid-like decrease. Interestingly, the response to electrical stimulation is different for each model, which underlines the relevance of using personalized models to guide the design of tES protocols.

## 4. Discussion

### 4.1. The role of chloride dynamics in ictal transitions

The present model sheds light on the potential role of pathological chloride accumulation and GABA depolarization in the generation of ictal activity in epilepsy. The addition of intracellular chloride dynamics to the NMM equations allows for the representation of spontaneous transitions from interictal to ictal state in an autonomous and realistic manner. This suggests that such transitions can be accounted for by the pathological accumulation of chloride inside pyramidal cells, as suggested by previous in vivo and in vitro studies [12, 49].

In the model, this pathological accumulation is generated by a dysfunction in the KCC2 co-transporter system, which has been extensively reported in epilepsy [12]. One of the model parameters, *α*_KCC2_, represents the degree of KCC2 transporter function in the simulated neuronal population, with higher values corresponding to a more functional transporter system. Increasing *α*_KCC2_ above a critical healthy threshold maintains the system in non-pathological background activity (see Appendix E) by avoiding the pathological Cl^−^ accumulation that leads to ictal activity. This points out to a crucial role of Cl^−^ transport pathology in the generation of seizures. Moreover, since extrasynaptic NMDA receptor activation induced by glutamate spillover is susceptible to activate the calcium-sensitive calpain that leads to KCC2 cleavage and downregulation [50], every seizure is susceptible to aggravate chloride homeostasis through the alteration of KCC2 expression in the membrane. Severe dowregulation of chloride transporter may lead to constant constitutive depolarizing GABA [16] whereas mild dowregulation could be associated with activity dependent overload of chloride in case of intense activity of GABAergic interneurons [8]. Both of these can be represented by the model through the *α*_KCC2_ parameter.

It is worth stating that the pathological expression of NKCC1 transporters in epileptic tissue, which has been also found in many cases [38, 51], could also exacerbate the high levels of chloride concentration in the pyramidal cells of the epileptogenic area. This mechanism has not been included in the model but is thought to have a complementary effect to the one of KCC2 transporters. In a future step, NKCC1-related mechanisms could be accounted for in the model equations.

A significant novelty of this model is the link between chloride accumulation in different locations of the pyramidal population and the amplitude of the PSPs generated upon GABA release. Previous models had exploited separately either the decrease of the amplitude of synaptic inhibitory gains [1] or the fact that GABA release may have a depolarizing effect in epileptic tissue [18]. In particular, the depolarizing effect of GABA release has been observed in the clinic and in animal models in numerous studies, many of which have pointed out to the potential implication of pathological chloride accumulation in this abnormal effect of GABAergic neurotransmitters [12, 8]. In our model, by simulating the pathological accumulation of Cl^−^ in the cell, we link the level of chloride concentration to the magnitude of GABAergic currents, and therefore with the amplitude of synaptic inhibitory gains, and from this we are able to simulate autonomous transitions from interictal to ictal activity — a novel advance in the field. This feature of the model captures the relationship between chloride transport dysfunction, GABA depolarization and epileptic seizures, and allows for the simulation of events in a seizure with other minimal assumptions, such as the presence of PV and SST interneuron populations.

We have proposed that the depolarizing effect of GABA in the seizure onset zone may be responsible for the initiation of seizures. At first sight, this seems to be in contradiction with the fact that some drugs such as Benzodiazepine (BZD), which are positive allosteric modulators of GABA_A_ receptors, are usually effective to prevent or to stop ongoing seizures in most patients suffering focal epilepsies [52]. Clinical cases for which BZD aggravate seizures exist, but remain uncommon [53]. However, it should be noted that BZD increases channel opening times and decreases the membrane resistance, prolonging shunting inhibition, and by this mechanism, retains its inhibitory effect, independently of the depolarizing or hyperpolarizing effect of GABA [54]. The mechanism of shunting inhibition has not been implemented in the current model, but see [18] for an example of shunting inhibition in a similar computational model.

### 4.2. Representation of depolarizing GABA physiology in the model can shed light on phenomenology of transitions

Several mechanisms have been proposed to explain the initiation of epileptic seizures, such as network excitation/inhibition imbalance [55], extracellular potassium concentration [56], or various channelopathies [57, 58]. Ours is an attempt to provide a succint, model-based insight into the potential pathophysiological mechanisms leading to seizure initiation. We have used as a starting point the discovery that excitation/inhibition imbalance in a simple NMM can reproduce the phenomenology observed in EEG and SEEG recordings during epileptic seizures [1], and we have proposed that this imbalance might be linked to the phenomenon of depolarizing GABA [8] by including pathological Cl^−^ dynamics in the model.

Indeed, the phenomenology of transitions in epilepsy is varied across patients, and even across seizures. The proposed model is able to capture some of the main features of these transitions, such as pre-ictal spikes, fast onset low voltage activity, or rhythmic activity. The parameter modifications required to represent some of these characteristics can be informative about the pathophysiological mechanisms responsible for epileptic seizures, as we now discuss.

Low voltage fast activity (LVFA) is usually found as a marker of seizure onset in epilepsy patients [59, 24, 44, 45]. This type of activity is generated in the model by a transition into a region of the parameter space where PV cells are active at very high frequencies. The transition to this region is due to decrease of IPSPs coming from SST cells, itself due to the pathological accumulation of chloride in the pyramidal cell dendrites. One should notice that the decrease of SST inhibition might also be due to the cell vulnerability or even cell death observed in several histological studies [60]. The model predicts that, next, the increased activity of somatic-targeting PV cells leads to the accumulation of chloride in the pyramidal cell soma. This in turn causes a decrease of GABAergic PSPs from PV interneurons, changing the system dynamics again, now from LVFA to rhythmic activity. Crucially, this evolution corresponds to the one presented in [1], where the transition to seizure was explained by a decrease in inhibition from dendritic-targeting slow interneurons first, then from perisomatic-targeting fast interneurons. Here we present a physiologically realistic explanation of this evolution, based on pathological chloride accumulation during seizure: first the dendritic compartments are more affected, then the somatic ones (see Figure 5).

Indeed, due to their low capacity, dendritic compartments are more susceptible to changes in chloride accumulation [12], and several studies have reported excitatory effects in dendritic GABA_A_ receptors when intensely activated [40, 61, 62]. Some studies have shown that optogenetic activation of PV or SST cells is equally effective in generating seizure-like activity [11], but others have reported that selective activation of perisomatic-targeting PV cells might be sufficient to trigger seizure-like activity [10, 9]. Although it is unclear whether the seizure is triggered in the first place by the depolarizing effect of GABA transmission from dendritic-targeting interneurons (as predicted by the model) or from somatic-targeting cells, in our model depolarizing GABA effects on both types of cells are needed to reproduce the whole interictal to ictal transition. Moreover, the mechanistic evolution presented might explain the patterns observed in epileptic seizure transitions, notably the presence of LVFA at seizure onset prior to clonic rhythmic activity.

Pre-ictal spikes appear in the model as a transition through a phase of spiking activity as dynamics evolve from background activity to fast onset activity. Chloride accumulation in the pyramidal cell population forces this change in the dynamics, which do not stabilize in the spiking phase region because the increased activity during this phase leads to more chloride accumulation in the pyramidal cells and thus, to a decrease of the GABAergic PSPs until the system enters a region of LVFA. This scenario is found in Patient 2 (see Figures 8(b) and 4(b)). Thus, if the transition from background to fast onset activity takes place in a more rapid manner, for example due to a very fast accumulation of chloride in the cell, pre-ictal spikes would be unlikely to appear, as the system would not spend enough time in the spiking activity region. This is the case for Patient 1 (see Figures 8(a) and 4(a)). Therefore, our results suggest that preictal spikes reflect a relatively slow transition through a region of spiking activity in parameter space, which exacerbates chloride accumulation and leads to the transition to either rhythmic activity or LVFA.

### 4.3. Modeling can characterize epileptogenic nodes in a patient-specific manner

We have provided a first methodology for the personalization of a node NMM from patient data. The model is capable of reproducing patient-specific, autonomous transitions to seizure-like activity that mimic some of the main features of the SEEG recordings in the most epileptogenic contact. This is a promising approach for future developments in computational modeling of epilepsy, since, to our knowledge, this is the first mesoscale modeling approach that by explicitly representing chloride regulation dynamics successfully reproduces transitions to ictal activity without the online manipulation of model parameters.

Previous attempts have relied on the heuristic tuning of one or several parameters for the same purpose [1, 18], providing a proof-of-concept of the relevance of an approach based on altered synaptic function. Building on this, in our model the initiation of the transition to seizure activity is triggered by a stochastic increase in the excitatory input received by pyramidal cells and changes in chloride dynamics. With respect to other, more abstract models that have also reproduced realistic seizures based on increases of excitation to the neural populations [1, 63], our model has the advantage that the additional equations that allow us reproducing such transitions represent physiological mechanisms—namely the chloride accumulation dynamics in pyramidal cells and the depolarizing GABA phenomenon in epilepsy.

A crucial result from the current study is the fact that the model has proven to be flexible enough to fit the wide variety of frequencies displayed in each phase of the transition from interictal to ictal activity in several patients. We have successfully fitted the model parameters for four different patients, each with different frequencies and features in each transition phase. This demonstrates the robustness of the model and of the personalization strategy presented, paving the way for the development of patient-specific computational models to guide invasive or non-invasive personalized treatments in epilepsy such as tDCS [64]. As an example, several studies have used neural masses or macroscale models embedded in a brain network to predict the outcome of surgery in epileptic patients [21, 20, 19, 65]. In these studies, the models of nodes placed in the epileptogenic zone are usually modified with respect to the rest of the network nodes with an increase of excitability, but they are rarely personalized based on the patient’s quantitative physiological data. The current model could be used in such network studies for a more realistic, personalized approach, where nodes in the epileptogenic zone display the phenomenology observed in physiological recordings. A network approach based on the current model is a natural next step, since our model is not only capable of reproducing seizure-like transitions, but also typical background or interictal activity (e.g., with healthy KCC2 transporters — high *α*_KCC2_ —, or with low excitatory input, see Appendix E).

Moreover, the personalized model is informative about several aspects of the patient’s pathology. The adjusted parameters, which are chosen to fit some specific features of SEEG recordings, represent realistic physiological variables, and, thus, reveal different characteristics of pathophysiology. Some examples include: the level of excitability of the epileptic tissue, represented by the value of *W*_*exc*_, the proportion of healthy cells in terms of baseline chloride accumulation (sometimes referred to as “static depolarizing GABA” [8], see Appendix B.3), given by the parameter *W*_*h*_, or the level of impairment of chloride transport, given by *α*_KCC2_. We leave for further work the analysis of these modeling aspects in larger, multi-patient datasets.

### 4.4. Physical modeling is needed to connect NMM dynamics with electrophysiology

The physics of SEEG measurements has a strong impact on the signals recorded. In particular, the morphology of SEEG signals is highly sensitive to the electrode location (see Figure 7). Previous attempts to generate realistic SEEG data with NMMs have used the average membrane potential perturbation in pyramidal cells as a surrogate for the electrophysiological signals generated by the neuronal population [1, 66, 18]. We have previously argued that this approximation might be inaccurate for the representation of realistic electrophysiological signals [22, 23]. Indeed, in the present work we have shown that the membrane potential signal is representative of the SEEG signal morphology at a very narrow range electrode depths, and, thus, cannot provide a full representation of the features of SEEG recordings. However, aspects of the signal such as the dominant frequencies in the different phases of the seizure are less influenced by the electrode position and can be captured by the average membrane potential perturbation in pyramidal cells. The spectrum of the SEEG signal is distorted across depths, but features such as frequency peaks are in general well preserved.

In view of this, and because in the clinical setting the imaging techniques used to retrieve the electrode position inside the brain are not precise enough for the purpose of SEEG morphology analysis (e.g., the resolution of computed tomography (CT) is around 0.5 mm), our personalization strategy is based on the spectral features of the signal, rather than in the morphology of the recordings (Figure 4). For the purpose of model personalization, we therefore use the average membrane potential perturbation in pyramidal cells, as was done in previous work.

Once the model has been personalized, however, a physical layer can be added to the NMM framework in order to achieve a more accurate representation of SEEG recordings. In recent work [67], we have provided methods to connect microscale neuron models with the neural mass formalism and with physical models of SEEG measurements, improving the accuracy of simulated electrophysiological signals. In the present study, we have used a simplified version of such framework (see Section 2.7), which includes a laminar version of the NMM embedded in a physical model. The electrode depth has been adjusted heuristically, based on the best fit with the real SEEG data. Future work could profit from more precise imaging techniques to retrieve the SEEG electrode position accurately, which would allow for the personalization of the combined NMM and physical model based on the recorded SEEG data.

### 4.5. The model can be used to represent the effects of brain stimulation in epileptic tissue

The aim of the computational model presented here is not only to provide a better understanding of the pathophysiology behind epileptic activity, but also to help designing personalized medical treatments for the disease. One promising direction for therapeutic interventions in epilepsy is transcranial electrical stimulation (tES), especially in cases where drugs and/or surgical intervention are not effective. The design of personalized stimulation protocols can be guided by computational models, which provide a unique tool to predict the outcome of electrical stimulation on the epileptic tissue. As a first step, by simulating the effect of electric fields in a personalized model that mimics the activity in the epileptogenic tissue of a given patient, it is possible to study in silico if the epileptic activity can be reduced, or suppressed, by tES targeting the epileptogenic zone.

The study presented in Section 3.4 constitutes an *in silico* proof of concept of the effects of tDCS in epileptogenic nodes, and shows how inhibition with external electric fields can reduce or even suppress the epileptiform activity in simulated nodes. Future work includes the analysis of the effects of weak electric fields at the network level and a more precise derivation of the coupling term *λ*_*n*_, which requires a translation of the effects of uniform weak electric fields from the microscale to the mesoscale. Work along these lines has been carried out in [68, 48], where the effects of electric fields are analyzed at the single neuron level. The emergence of exact mean field theories deriving neural mass models from microscale ones, the construction of detailed population models, as well as experimental work can all help shed some light on the proper rescaling of the coupling parameter in our models [69].

## 5. Conclusion

It has become apparent during the last decades that computational models of epilepsy can provide key insights for treatment of neurological diseases like epilepsy. In this work we have provided a neural mass model combining physiology and biophysics to simulate electrophysiological signals from SEEG recordings in patients. By including a dynamical mechanism for GABA depolarization, we have developed an autonomous model of realistic interictal to ictal transitions. The model can be personalized at the single epileptic node level from SEEG recordings by including also physical aspects associated with the measurement process in SEEG: to better represent electrophysiological measurements, the model captures the laminar architecture of the neocortex and can be embedded in a realistic physical head model. The development of this model paves the way for robust personalization methods in epilepsy. We have demonstrated personalization in this modeling framework analyzing SEEG data from four epilepsy patients.

## Acknowledgments

E.L.-S., R.S.-T., F.B., F.W., P.B. and G.R. designed research; E.L.-S., R.S.-T., E.L., M.G., P.B and G.R. performed research; R.S.-T., E.L., E.K.-E., M.Y., B.M., R.S., J.Mo. contributed new reagents or analytic tools; E.L.-S., E.L., J.Ma. and D.L.-S. analyzed data; E.L.-S. and G.R. wrote the paper.

E.L.-S., R.S.-T., E.L., B.M., R.S. and D.L.-S. work for Neuroelectrics Barcelona. G.R. is co-founder of Neuroelectrics.

All the authors are partially supported by the ERC Synergy grant Galvani. This work has received funding from the European Research Council (ERC) under the European Union’s Horizon 2020 research and innovation programme (grant agreement No 855109).

## Ethics statement

This study was approved by the institutional review board of the Assistance Publique Hopitaux de Marseille, and informed written consent was obtained for all patients.

## Appendix A. SEEG data analysis

### Appendix A.1. Peak frequency estimation

Separate time-frequency representations (TFR) of oscillatory power were carried out for slow and fast frequencies [26]. For slow frequencies (0.5–30 Hz) the frequency axis was estimated at 30 logarithmically spaced frequencies using a fixed sliding time window of 2 seconds duration. Prior to the FFT, each time-window was multiplied by a single Slepian taper resulting in a frequency smoothing of 1 Hz. For high frequencies (30–200 Hz), the frequency axis was estimated at 60 logarithmically spaced frequencies using the same sliding window duration as for the slow frequencies. Prior to the FFT, time windows were multiplied by three Slepian tapers resulting in a 3 Hz frequency smoothing. The power of each TFR was normalized to decibels (dB) relative to the maximum value within the entire time and frequency matrix.

The TFR for high and low frequencies allowed us to easily identify the fast onset and the rhythmic clonic phases, respectively. For each patient and seizure, we manually defined the onset and offset of the two events and we automatically estimated their duration and peak frequency. The peak frequency of the SEEG signal recorded in the most epileptogenic node was obtained by spectral parametrization following the approach described in [27]. This method assumes that the power spectrum density *PSD* can be reconstructed by the sum of *N* Gaussian functions *G*_*n*_ plus a Lorentzian function *L*:

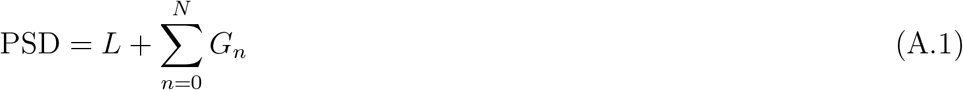

Briefly, the algorithm flattens the PSD by subtracting a Lorentzian fit. Then, an iterative process identifies peaks in the flattened PSD by fitting Gaussian functions. These Gaussians are subtracted from the original PSD to fit again a Lorentzian to improve its estimate. Finally, the goodness of fit is computed by adding the *L* and *G*_*n*_ and comparing it to the raw PSD (see [27] for more details). The Gaussian function is defined by

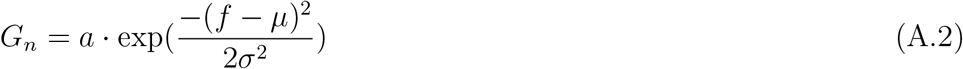

where *a* is the power of the peak (decibels), *µ* is the center frequency (Hz), *σ* is the standard deviation of the Gaussian (Hz) and *f* is the vector of input frequencies (Hz). The Lorentzian function is defined as

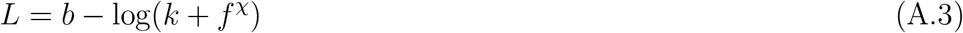

where *b* is the broadband offset, *χ* is the exponent and *k* is the parameter that controls the bend of the aperiodic component [70, 71]. To parametrize the fast onset, we set k=0 [71] and we fit up to three Gaussians to capture its prototypical chirp dynamics [44]. We calculated the average *µ* taken over the three Gaussians (*µ*_*c*_), and then we computed the average over seizures for each patient; this average fast onset frequency was used for the creation of the personalized model. For slow frequencies, the same procedure was used, with the only exception that we fitted the rhythmic tonic dynamics with *k* ≠ 0 to account for the fact that the PSD knee [27] for slow frequencies appeared around 10 Hz.

### Appendix A.2. Time-frequency representation computation

In order to capture the high frequency traces of the low-voltage fast activity (LVFA) and the rhythmic activity of the ictal phase in the same picture, we have computed weighted time-frequency representations of the SEEG signals (see Figures 5 and 8). The frequency axis was estimated at 2 Hz linearly spaced frequencies using a fixed sliding time window of 0.5 seconds. For each time window the frequency values are normalized, so that a maximum power value per time point is shown in the time-frequency representation.

## Appendix B. Summary of the extended neural mass model

Our extended NMM is characterized by state variables describing the firing rate of each neuronal population, the average membrane perturbations induced by each synapse, and the average chloride concentration in each pathological population synapse. The activity of the NMM is then mapped into physical measurement space (see next section). We provide the detailed NMM equations below.

### Appendix B.1. Firing rate and synaptic equations

The population state *𝒫*_*n*_ is first characterized by the average membrane potential *v*_*n*_ of the cells in the population — itself the summation of all the average pre-synaptic membrane perturbations *u*_*s*_ — and by its average firing rate *φ*_*n*_, which is computed using a non-linear function of the membrane potential,

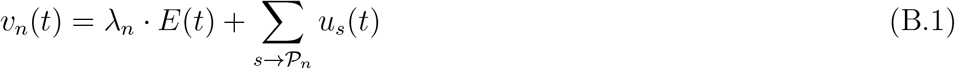

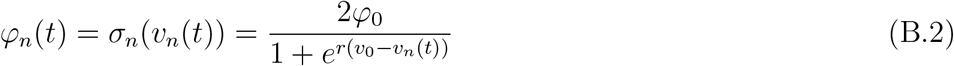

where the sum in the first equation is over synapses reaching the population, *φ*_0_ is half of the maximum firing rate of each neuronal population, *v*_0_ is the value of the potential when the firing rate is *φ*_0_ and *r* determines the slope of the sigmoid function associated to the neuronal population, *σ*_*n*_, at the central symmetry point (*v*_0_, *φ*_0_) [6, 5, 7].

The term *λ*_*E*_ · *E*(*t*) represents the average membrane perturbation induced by an external electric field [31, 32] on the population and accounts for the effects of electrical stimulation or ephaptic effects [34], in the case where they are to be included (see 3.4).

Each synapse *s* is described by an equation representing the conversion from an input firing rate *φ*_*n*_ from the pre-synaptic population *n* into an alteration of the membrane potential *u*_*s*_ of the post-synaptic neuron. This relation is represented by the integral operator 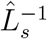 (a linear temporal filter), the inverse of which is a differential operator 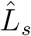,

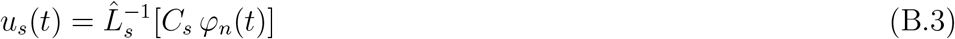

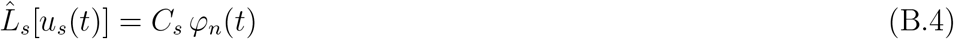

where *C*_*s*_ is the connectivity constant between the populations. The differential operator 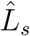 is defined as

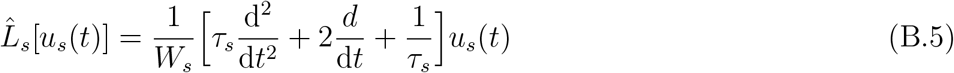

where *W*_*s*_ is the average excitatory/inhibitory synaptic gain and *τ*_*s*_ the synaptic time constant. This time constant is a lumped parameter representing the population averaged effective delay and filtering time associated with the time from reception of input in the cell to soma potential perturbation (it is not, e.g., the time constant of the ion channel or even the local synaptic time constant at the dendrite).

To motivate these equations, we note that to find the solution to Equation B.4, one can first solve

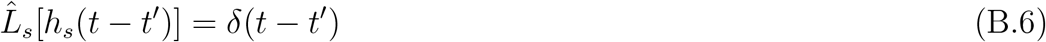

with the boundary conditions a) *h*_*s*_(*t*′) = 0 for *t* ≤ 0, and b) requiring a finite discontinuity of the first derivative at *t* = *t*′. The solution is then obtained from the properties of the Dirac delta function (∫ *dt*′ *f*(*t*′)*δ*(*t*′ − *t*) = *f*(*t*)),

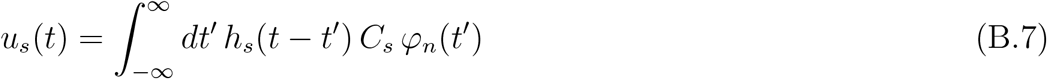

The function *h*_*s*_(*t*−*t*′) is known in mathematics as the Green’s function of the operator, and as the PSP (post-synaptic potential) in the neural mass modeling community (see, e.g., [7]). It is is given by

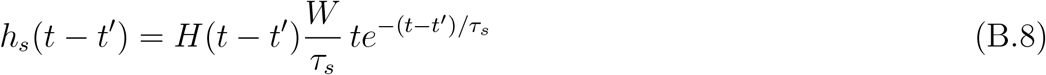

with *H*(*t* − *t*′) the Heaviside function (equal to 0 for *t* < *t*′, 1 otherwise). We note in passing that the integral of *h*_*s*_(*t* − *t*′) is given by 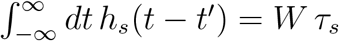.

The detailed equations for each synapse that follow in the model described in the main text are (for the case where no external electric field or ephaptic effects are considered):

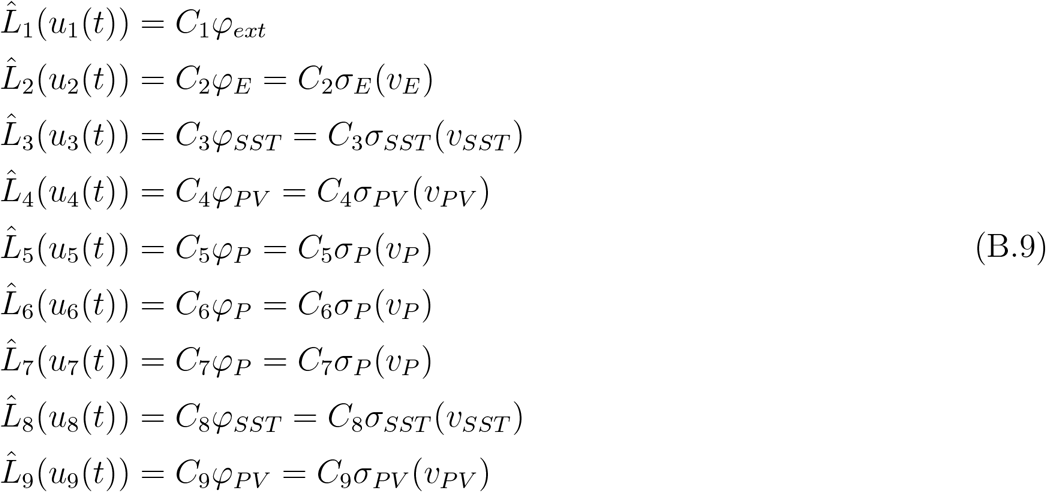

with neuron membrane potentials given by

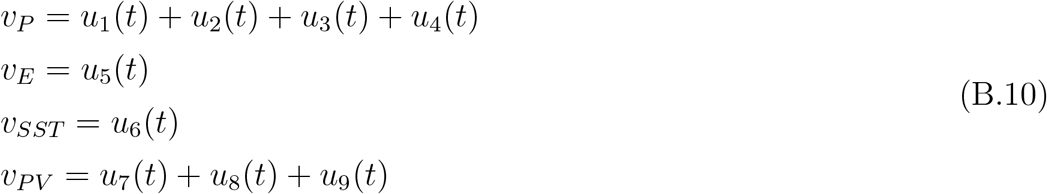

The pre- and post-synaptic populations corresponding to each synapse index are:

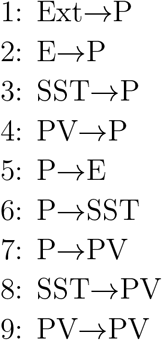

A description of the neural mass model parameters is given in Table B1, and a schematic summary of the model equations (including the chloride dynamics model described in the next section) is provided in Figure 3.

**Table B1:** Description of the neuron and synapse parameters. Note that the first three parameters (*W*_*s*_, 1*/τ*_*s*_, *C*_*s*_) are synapse-specific, the sigmoid parameters (*v*_0_, *φ*_0_, *r*) are common to all neuronal populations

| Parameter | Description |
| --- | --- |
| $W_s$ | Average excitatory and inhibitory synaptic gain |
| $1/\tau_s$ | Time rate constant of average excitatory and inhibitory post synaptic potentials |
| $C_s$ | Average number of synaptic contacts between population types |
| $v_0$ | Potential when 50% of the firing rate is achieved |
| $\varphi_0$ | Half of the maximum firing rate |
| $r$ | Slope of the sigmoid function at $v_0$ |
| $p_m, p_{std}$ | Mean and standard deviation of the external input to the pyramidal population |

### Appendix B.2. Chloride dynamics equations

The concentration of Cl^−^ in a synaptic location is calculated according to the balance equations as a function of the combined extrusion of chloride by KCC2 transporters and the firing-rate-dependent influx of chloride in GABA_A_ synapses, modulated by a surface-to-volume translating parameter 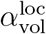 that depends on the synapse location (apical dendrites or perisomatic region),

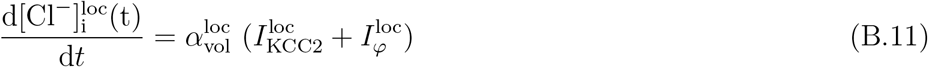

The Cl^−^ flux associated with KCC2 transporters is given by

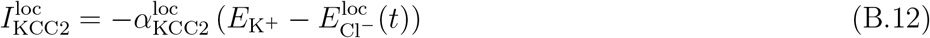

with

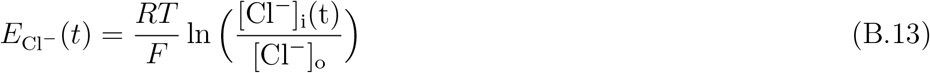

We assume here that the reversal potential of potassium 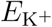 remains constant, as does the membrane potential *V*_m_. The parameter *α*_KCC2_ reflects the rate of extrusion of Cl^−^ through KCC2 transporters. The Cl^−^ current through GABA_A_ channels due to the activity of inhibitory interneurons is given by

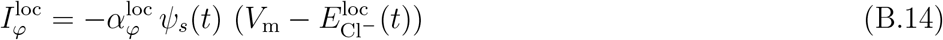

with

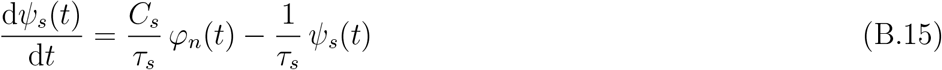

The parameter 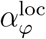 relates the activity of GABAergic interneurons with the Cl^−^ currents. This parameter can be personalized on a patient-specific basis.

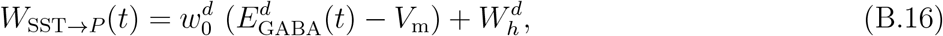

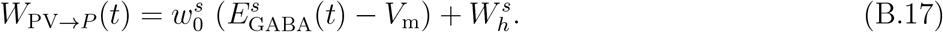

with

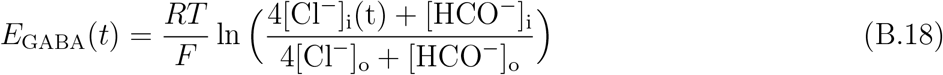

Other synaptic gains (pyramidal population to interneurons, SST to PV, etc.) are assumed to be constant in the model. We assumed as well that all ionic concentrations are constant to a good approximation except for the concentration of chloride inside the cells [Cl^−^]_i_.

A description of the chloride dynamics parameters used in the model is provided in Table B2, and an illustrative diagram of the model equations is shown in Figure 3.

### Appendix B.3. Chloride-dependent synaptic gain dynamics. Contribution from sub-populations with static gains

In the NMM framework, synaptic gains (*W*) represent the mean amplitude of the PSP generated by a given synapse type in a cell population. Synaptic gains of GABAergic synapses are proportional to the current generated in those synapses,

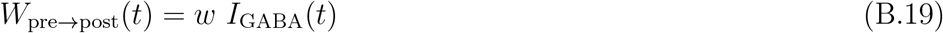

where *w* is the proportionality factor linking PSP amplitude to synaptic current, and *I*_GABA_ is the GABAergic synaptic current, given by

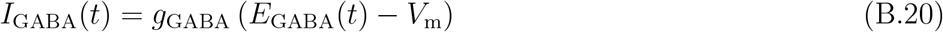

where *g*_GABA_ is the average inhibitory conductance, *V*_m_ is the membrane potential (assumed approximately constant) and *E*_GABA_ is the reversal potential of GABA. Thus, we can write

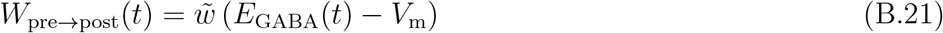

**Table B2:**
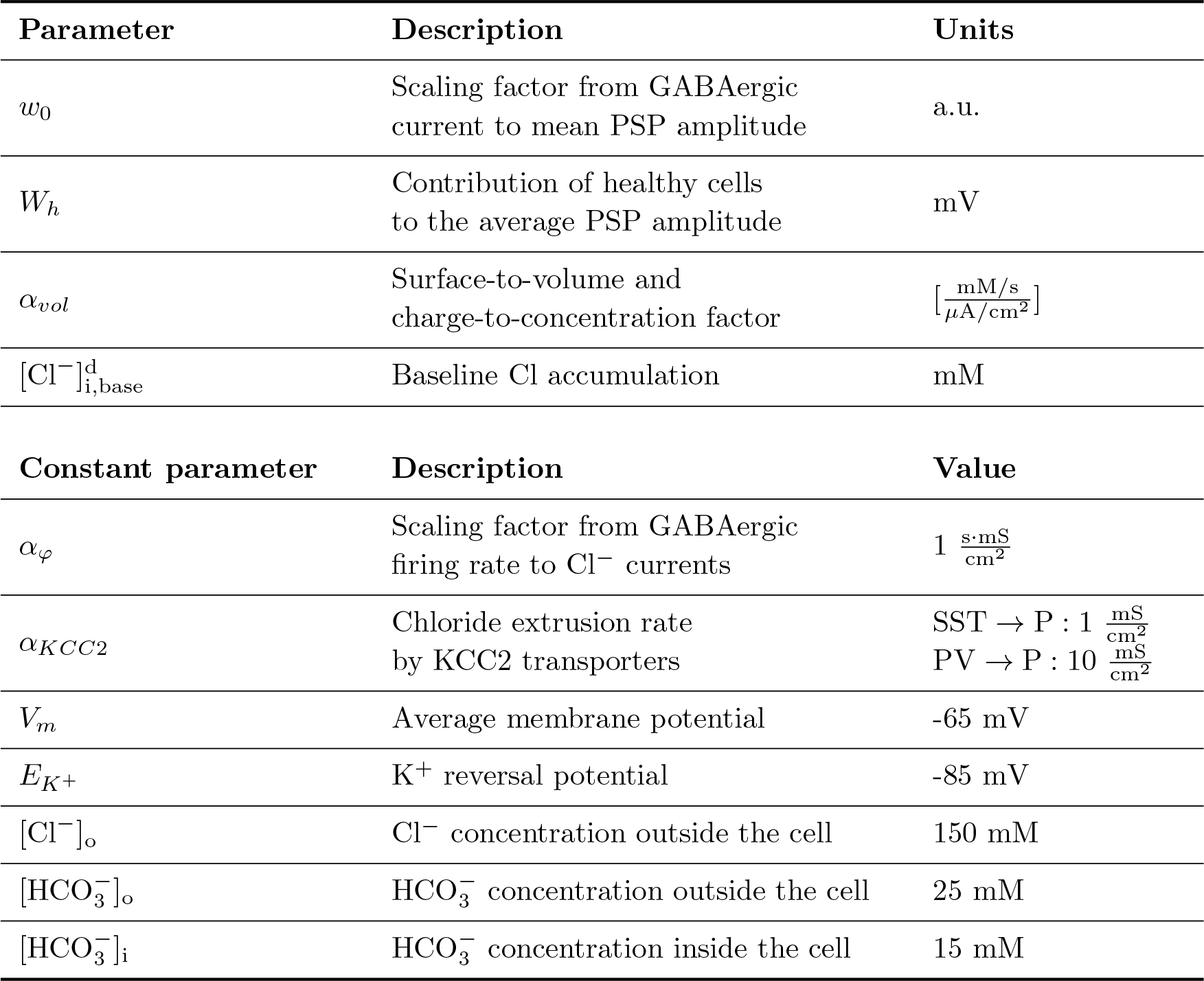
Parameters, description and values of the Cl^−^ dynamics parameters.

| Parameter | Description | Units |
| --- | --- | --- |
| $w_0$ | Scaling factor from GABAergic current to mean PSP amplitude | a.u. |
| $W_h$ | Contribution of healthy cells to the average PSP amplitude | mV |
| $\alpha_{vol}$ | Surface-to-volume and charge-to-concentration factor | $\left[ \frac{\text{mM/s}}{\mu\text{A/cm}^2} \right]$ |
| $[\text{Cl}^-]_{i,\text{base}}^d$ | Baseline Cl accumulation | mM |
| Constant parameter | Description | Value |
| $\alpha_\varphi$ | Scaling factor from GABAergic firing rate to $\text{Cl}^-$ currents | $1 \frac{\text{s}\cdot\text{mS}}{\text{cm}^2}$ |
| $\alpha_{KCC2}$ | Chloride extrusion rate by KCC2 transporters | SST → P : $1 \frac{\text{mS}}{\text{cm}^2}$<br>PV → P : $10 \frac{\text{mS}}{\text{cm}^2}$ |
| $V_m$ | Average membrane potential | -65 mV |
| $E_{K^+}$ | $\text{K}^+$ reversal potential | -85 mV |
| $[\text{Cl}^-]_o$ | $\text{Cl}^-$ concentration outside the cell | 150 mM |
| $[\text{HCO}_3^-]_o$ | $\text{HCO}_3^-$ concentration outside the cell | 25 mM |
| $[\text{HCO}_3^-]_i$ | $\text{HCO}_3^-$ concentration inside the cell | 15 mM |

with 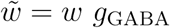.

We slightly extend the above to assume that in the pyramidal cell population represented by the NMM there are sub-populations with different properties. For example, consider the case where there is a proportion *α*_*h*_ of healthy neurons where the transport of chloride is not dysfunctional. In those neurons, the concentration of chloride inside the cell is kept constant by the Cl^−^ transport system, and, thus, the reversal potential of GABA is *static*. Let *E*_*GABA,h*_ be the reversal potential of GABA for the healthy population. Then, the average gain representing two sup-populations of healthy and pathological cells can be written as

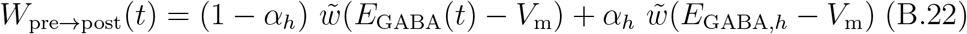

where the first term represents the chloride-dependent pathological variation of synaptic gains, and the second a static contribution from healthy cells to the average PSP.

We then define 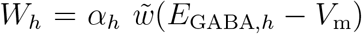 and 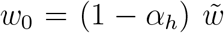 to bring us to the formulation used in the main text,

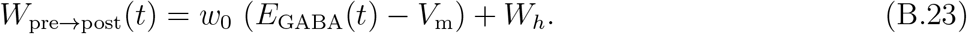

For the specific synapses of the model with pathological chloride accumulation, this leads to Equations 13 and 14 in the main body,

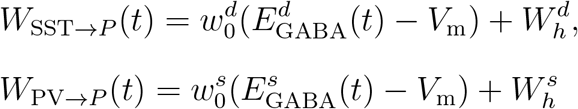

Similarly, consider the case where there is another sub-population of cells where the KCC2 transport system is weakened to the point where the chloride concentration is the same inside and outside the cell. Because the GABA reversal potential is affected by the concentration of chloride and 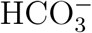, in this situation *E*_GABA_ is again a static quantity, but it can lead to a static positive value of *W*. To see this, recall that *E*_*GABA*_ is the reversal potential of GABA (the joint reversal potential of Cl^−^ and 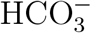) and can be calculated using the Goldman-Hodgkin-Katz (GHK) equation,

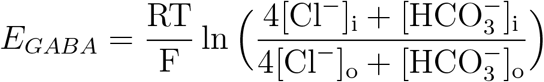

Consider as healthy values [Cl^−^]_i_ = 6 mM, [Cl^−^]_o_ = 150 mM, and 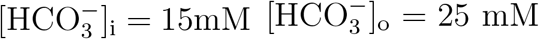. With RT*/*F = 25.693 mV, we find *E*_*GABA*_ = −71.3 mV. If instead we take the extreme case where the concentration of chloride is the same inside and outside the cell, [Cl^−^]_i_ = 150 mM, we find *E*_*GABA*_ = −0.4 mV, leading to a depolarizing effect of the synapse (recall that the resting potential *V*_m_ is about −65 mV). Thus, in this case the static gain contribution can represent pathological sub-populations where the GABA reversal potential is static but in an unhealthy regime (static depolarizing GABA [8]).

In summary, the static component *W*_*h*_ represents the aggregate effects of static intracellular chloride ionic concentration on GABA gain from existing sub-populations (healthy or unhealthy).

## Appendix C. Physical model for generation of SEEG recordings

### Appendix C.1. NMM-derived voltage

There is a wide agreement that the extracellular voltage recordings in the low frequency range are to a large extent generated by membrane currents that appear due to synaptic inputs in pyramidal cells. This is due to the coherence in space and time [72] of these currents that allows them to contribute in an additive manner into the extracellular voltage.

When neurotransmitters act on synaptic receptors a membrane current appears. An excitatory synapse produces an inward current that is seen as a negative current source (i.e. a sink) from the extracellular medium. Conversely, an inhibitory synapse causes an outward current that is seen as a positive current source (i.e. a source) from the outside. Within the timescale of SEEG recordings, these current sinks or sources are always balanced by a passive return current to achieve electroneutrality. The synaptic currents and their associated return currents can give raise to sizable dipoles (or higher order n-poles) and if many of these appear synchronized in time, a measurable voltage perturbation is produced.

According to compartmental models of neurons, the distribution of sources and sinks that appears due to a synaptic input depends on the location of the synapse [73]. For example, a synapse on the apical dendrites in the layer 1 of a pyramidal cell generates a return current that is mostly clustered around the soma. Interestingly, a synapse on the basal dendrites generates a return current that is mostly localized around the soma as well [74]. Thus, return currents can be either localized near the synapse injection site or they can appear in remote locations.

In this study, we built a simplistic model to simulate the voltage recorded by SEEG contacts and connect the NMM framework with real physical measurements. To do so, we simulated the voltage generated by layer 5 pyramidal cells only and we considered two possible synaptic locations: layer 1 and layer 5. A simplistic model, consisting in discrete sources, was used to approximate the depth profile of the current source density produced by a synaptic input in those two cases (see Figure C1). For simplicity, these discrete sources were located at depths corresponding to approximately the cortical layer depths. Namely, an excitatory synaptic input in layer 1 was modeled with a discrete current source of strength −*I* located there and two other sources of strength 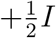 (the return current) located at layers 4 and 5. An excitatory synaptic input in layer 5 was modeled with a discrete source of strength −*I* located there and a return current −*I* located in layer 4. Inhibitory inputs were modeled with the exact same distribution but with opposite signs.

### Appendix C.2. Physical model

The current source model described above was used to generate a model of a pair of consecutive contacts in an SEEG lead embedded in a cortical patch. The model consisted of 3 tissue layers to represent the white matter (WM), the gray matter (GM) and the cerebrospinal fluid (CSF). The dimensions of the model and the conductivity assigned to each layer can be found in Figure C2(a). We assumed an SEEG lead completely perpendicular to the cortical surface which allowed building an axisymmetric model. SEEG contacts were modeled as 0.8 mm diameter and 2 mm length cylinders separated by an insulating part with 1.5 mm of length. A conductivity of 1000 S/m was set to the contacts and of 10^−5^ S/m to the insulating part between them. The current sources were modeled as arrays of discrete point sources (i.e. line current sources given that the model is axisymmetric) evenly distributed every 100 *µ*m (see Figure C2(a)). These arrays were placed at three different depths from the cortical surface (0.25, 1.45 and 1.85 mm) corresponding to the cortical layers 1, 4 and 5 and they extended up to a radial distance of 2 cm from the electrode. A radial distance of 100 *µ*m was left between the SEEG contacts and the nearest source to account for tissue damage due to electrode insertion.

**Figure C1:**
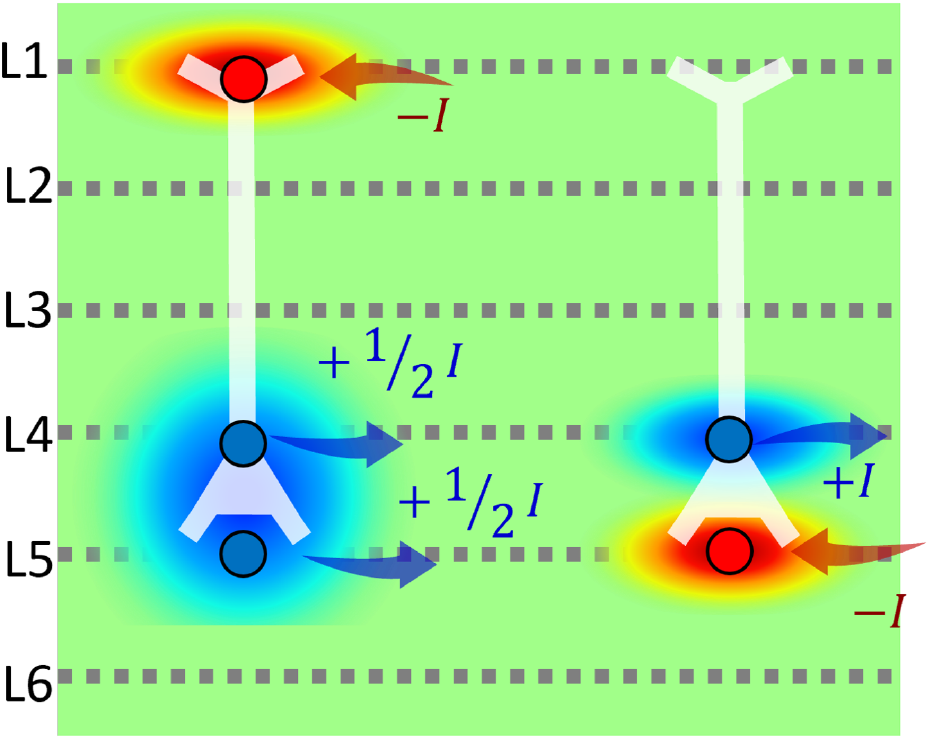
Current distributions used to model a synapse at layer 1 (left) and a synapse at layer 5 (right) in a layer 5 pyramidal cell.

The electric potential distribution was calculated by solving Laplace equation:

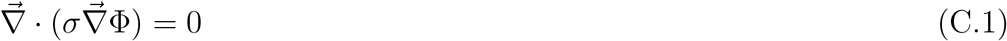

With the additional boundary conditions of the current sources and a floating potential with zero current assigned to the contact surfaces.

The problem described above was solved using COMSOL Multiphysics 5.3 (Stockholm, Sweden) for different depths of the SEEG electrode and for the two possible synaptic locations (with their associated current distribution models). These solutions were then used to generate SEEG recordings from NMM results.

For a given depth of the SEEG contacts, *z*_*el*_ the voltage difference over time between them can be calculated as a weighted sum of the aggregated post-synaptic current of each synapse *I*_*s*_:

**Figure C2:**
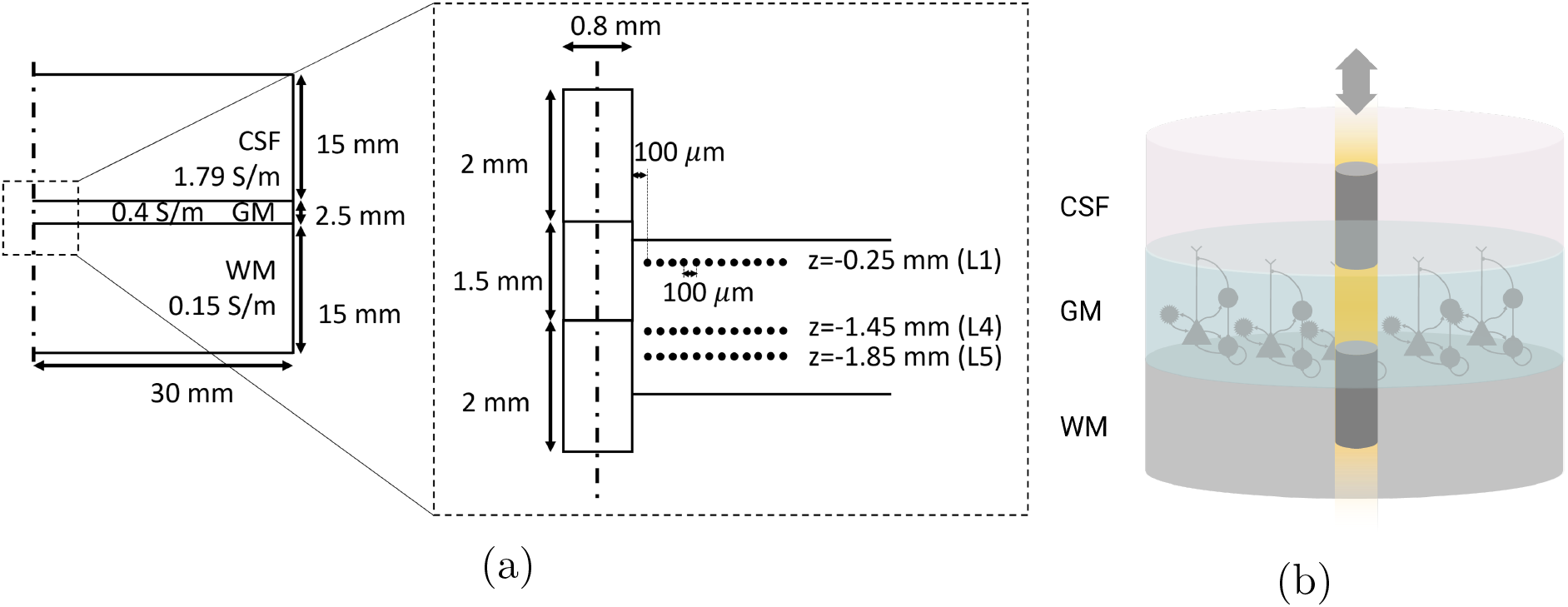
a) Schematic of the 3 layer FEM axisymmetric model used to simulate the voltage difference between a pair of SEEG contacts. b) Illustrative 3D view of the physical model. Conductivity values are taken from [43].

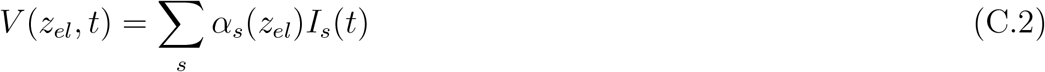

where *α*_*s*_ (in V/A) represents the contribution of each synaptic current to the measured voltage and depends on the electrode depth. In this study we only considered two types of post-synaptic currents (layer 1 and layer 5). The contribution of each of them was extracted from the solutions of the FEM described above. Namely, each coefficient *α*_*s*_ was calculated as the voltage difference generated by a synaptic unit current *I* in every point of the array with the corresponding depth distribution.

The post synaptic currents over time, *I*_*s*_(*t*), are computed from the output of the NMM as

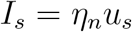

*η*_*n*_ is a conversion factor that relates the average potential perturbation to a physical synaptic current in the receiving population. Each synapse perturbation at the single neuron level will produce a local flow of ions across the membrane and, therefore, a local micro-scale synaptic current. Therefore, we assume that the population membrane perturbation *u*_*s*_ is related to the aggregated synaptic current *I*_*s*_ through the proportionality gain factor *η*_*n*_, which depends on the neuron population characteristics, such as cell density and cell morphology. To generate SEEG signals with realistic magnitudes, we have chosen *η*_*n*_ = 10^−8^ A/mV.

## Appendix D. Comparison of real and simulated SEEG signals

**Table D1:**
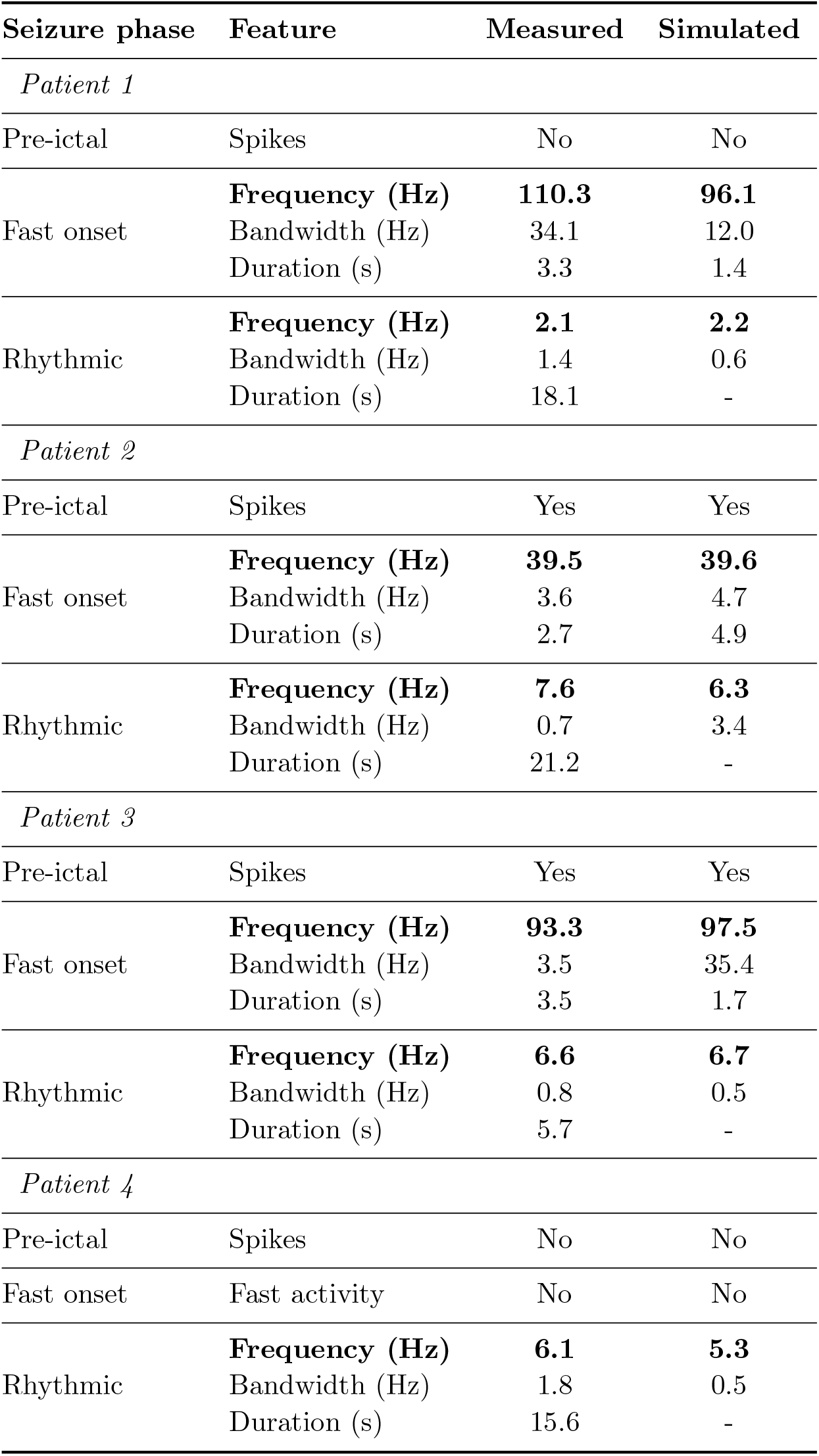
Main features of the SEEG signal recorded at the contact positioned in the most epileptogenic area of each patient (average over all seizures) and comparison with the simulated SEEG signal. The duration of the rhythmic phase in simulated data is not shown because no seizure termination mechanisms are included in the model (see *Methods*). The time intervals selected for the computation of the representative frequencies are shown in Figure 8.

| Seizure phase | Feature | Measured | Simulated |
| --- | --- | --- | --- |
| *Patient 1* |  |  |  |
| Pre-ictal | Spikes | No | No |
| Fast onset | **Frequency (Hz)** | **110.3** | **96.1** |
|  | Bandwidth (Hz) | 34.1 | 12.0 |
|  | Duration (s) | 3.3 | 1.4 |
| Rhythmic | **Frequency (Hz)** | **2.1** | **2.2** |
|  | Bandwidth (Hz) | 1.4 | 0.6 |
|  | Duration (s) | 18.1 | - |
| *Patient 2* |  |  |  |
| Pre-ictal | Spikes | Yes | Yes |
| Fast onset | **Frequency (Hz)** | **39.5** | **39.6** |
|  | Bandwidth (Hz) | 3.6 | 4.7 |
|  | Duration (s) | 2.7 | 4.9 |
| Rhythmic | **Frequency (Hz)** | **7.6** | **6.3** |
|  | Bandwidth (Hz) | 0.7 | 3.4 |
|  | Duration (s) | 21.2 | - |
| *Patient 3* |  |  |  |
| Pre-ictal | Spikes | Yes | Yes |
| Fast onset | **Frequency (Hz)** | **93.3** | **97.5** |
|  | Bandwidth (Hz) | 3.5 | 35.4 |
|  | Duration (s) | 3.5 | 1.7 |
| Rhythmic | **Frequency (Hz)** | **6.6** | **6.7** |
|  | Bandwidth (Hz) | 0.8 | 0.5 |
|  | Duration (s) | 5.7 | - |
| *Patient 4* |  |  |  |
| Pre-ictal | Spikes | No | No |
| Fast onset | Fast activity | No | No |
| Rhythmic | **Frequency (Hz)** | **6.1** | **5.3** |
|  | Bandwidth (Hz) | 1.8 | 0.5 |
|  | Duration (s) | 15.6 | - |

**Figure D1:**
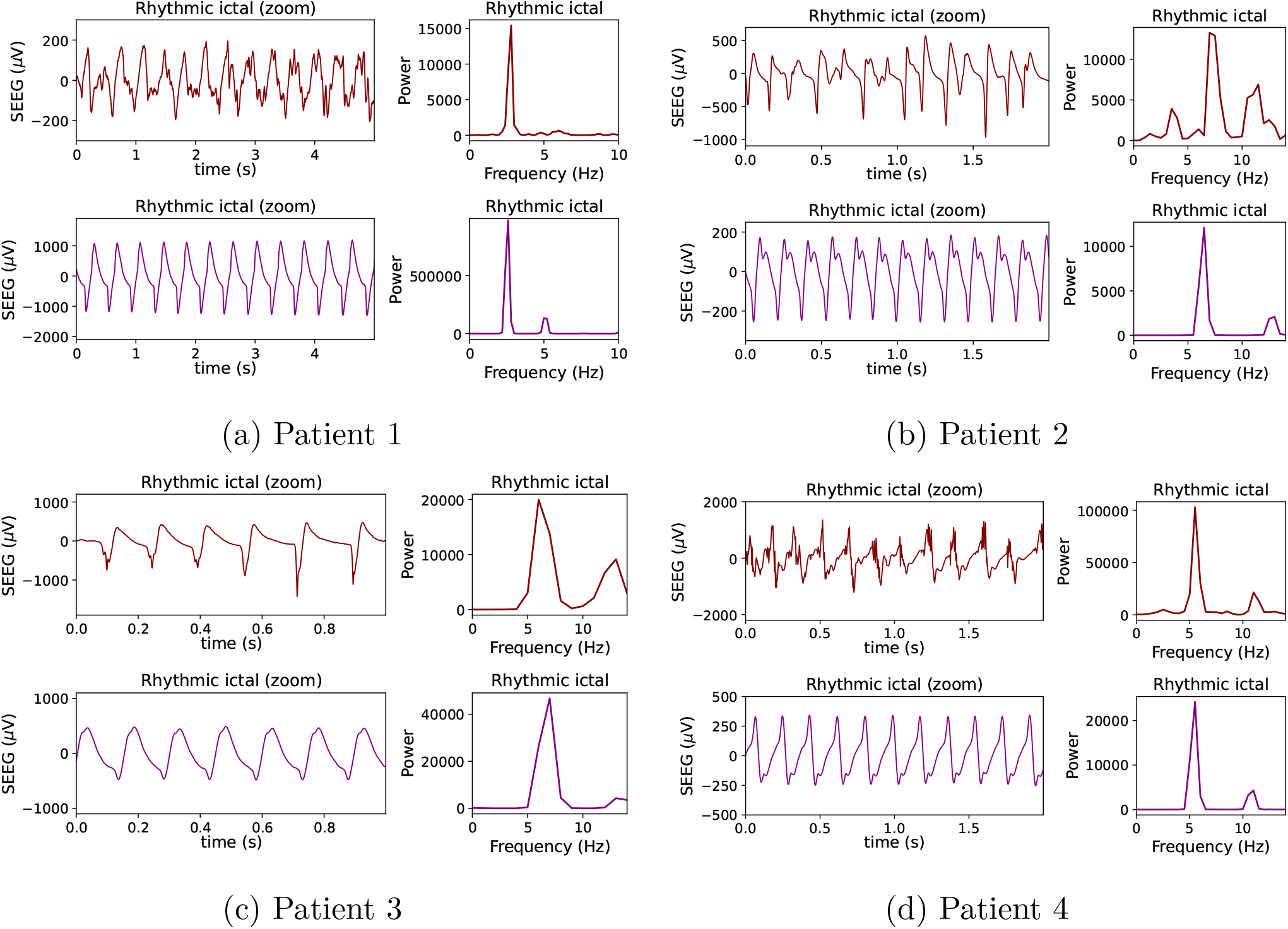
Detailed morphology of the rhythmic ictal phase signal for real (maroon) and simulated (purple) SEEG data. For each patient, the panels in the left represent the real and simulated SEEG data, and the right panels show the PSD of the signal. The simulated SEEG signal corresponds to a particular location of the SEEG electrode in the physical model, determined by its depth inside the grey matter *z*_*el*_, which provides a reasonable fit of the signal morphological features. Patient 1: *z*_*el*_ = −1.6 mm; Patient 2: *z*_*el*_ = −1.55 mm ; Patient 3: *z*_*el*_ = −1.6 mm; Patient 4: *z*_*el*_ = −6.05 mm.

**Figure D2:**
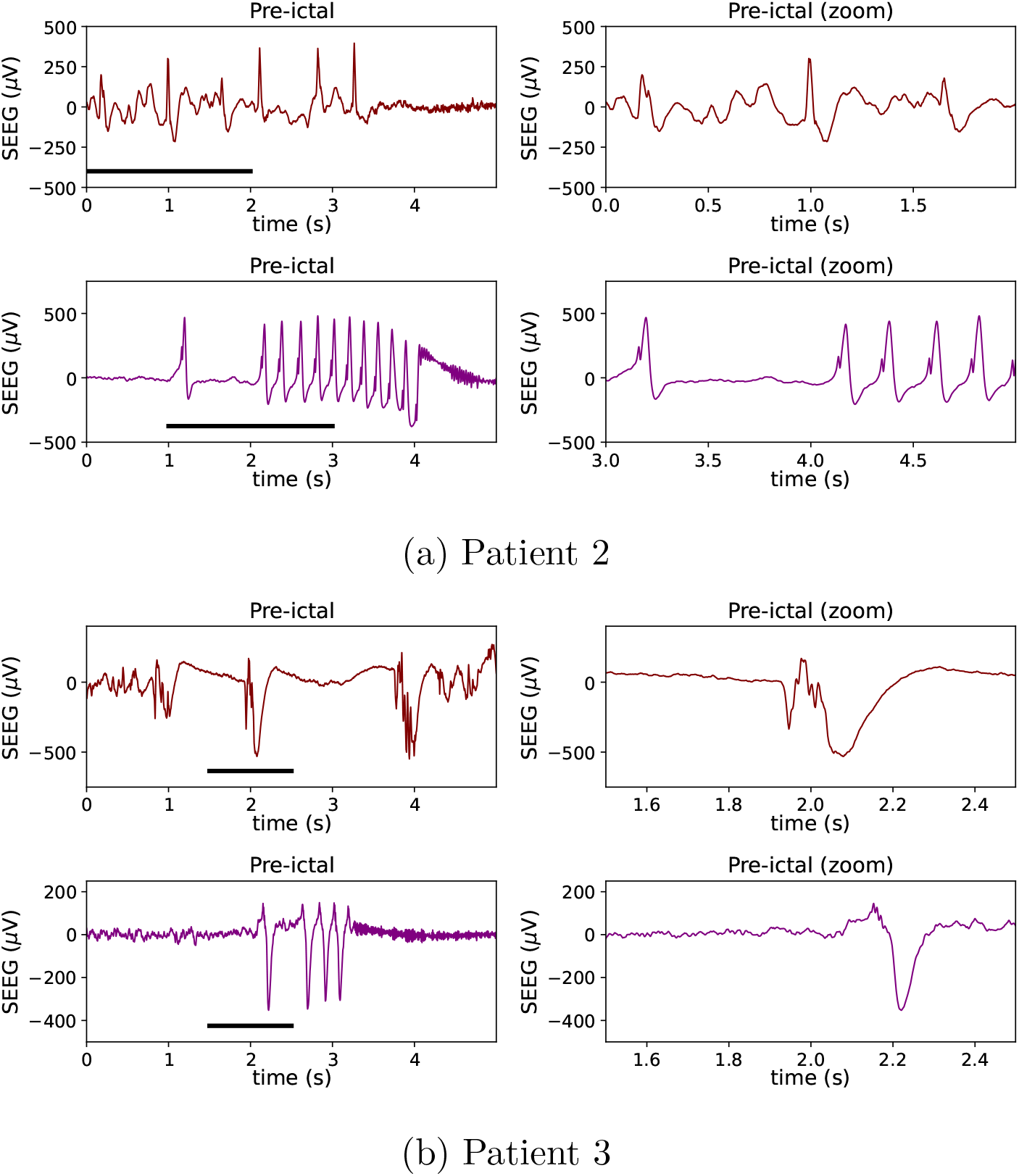
Detailed morphology of the pre-ictal phase signal for real (maroon) and simulated (purple) SEEG data. For each patient, the panels in the left represent the real and simulated SEEG data, and the right panels show a zoomed view of the signal (the time period selected is marked with a black trace in the left panel). The simulated SEEG signal corresponds to a particular location of the SEEG electrode in the physical model, determined by its depth inside the grey matter *z*_*el*_, which provides a reasonable fit of the signal morphological features. Patient 2: *z*_*el*_ = −6.0 mm; Patient 3: *z*_*el*_ = −6.05 mm.

## Appendix E. Non-pathological models

We have compared the personalized model for Patient 2 with two non-pathological versions of the model. In the first non-pathological version of the model, we have doubled the value of the parameter *α*_KCC2_ (in the soma and in the dendrites, see Table 2 of the main text for the original values). This corresponds to a situation where more KCC2 transporters are functional (twice as much). As predicted, the chloride accumulation reaches lower values than in the pathological model both in the dendrites and the soma, and the system does not enter into seizure (see Figure E1).

In the second version, we decreased the average value of the external input from 90 s^−1^ to 45 s^−1^. This variation corresponds to a decrease of the excitation to the NMM. It can be seen that the transition to ictal state is avoided, and the values of chloride accumulation stabilize at lower values than in the pathological model.

**Figure E1:**
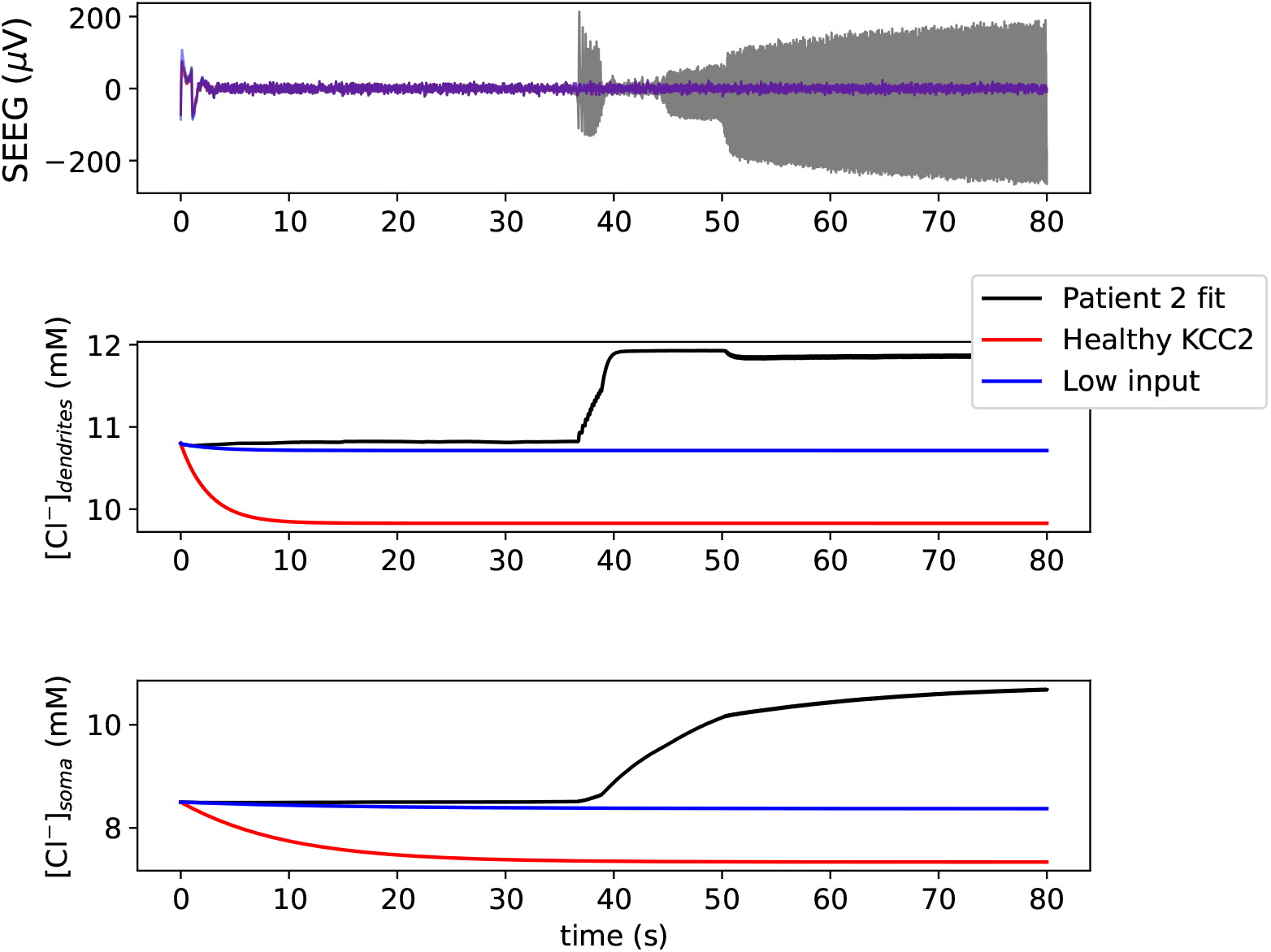
Comparison between the personalized model for Patient 2 (grey traces) with two non-pathological versions of the model (red and blue traces). The top panel displays the simulated SEEG signal (for an electrode depth *z*_*el*_ = −1.55 mm); the middle and bottom panels represent the chloride accumulation in the dendrites and the soma, respectively. Red traces correspond to increased function of KCC2 transporters and blue traces to decreased external input to pyramidal cells. In both cases the transition to ictal state is avoided.

## Supplementary Information

**Fig. S1.**
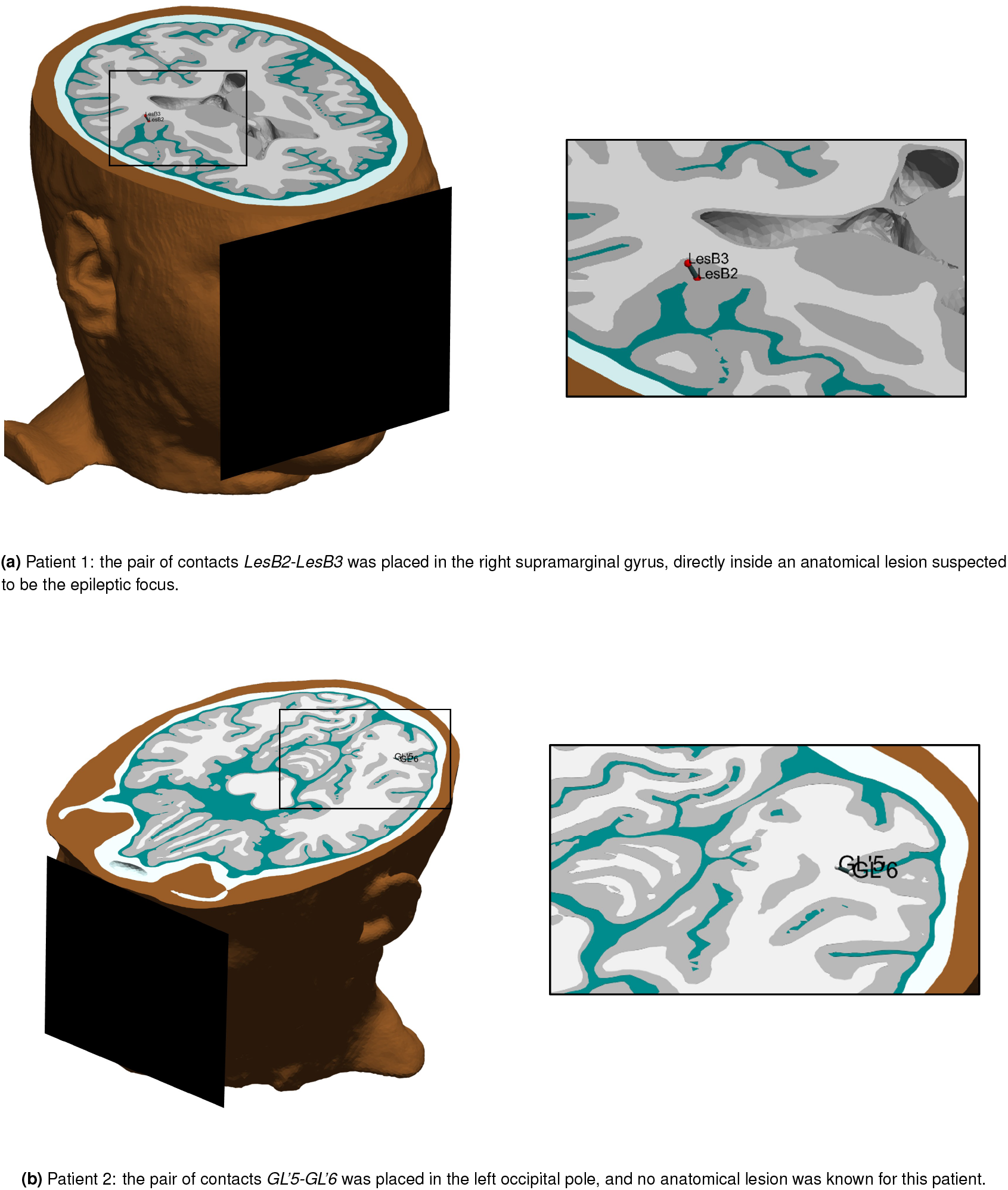
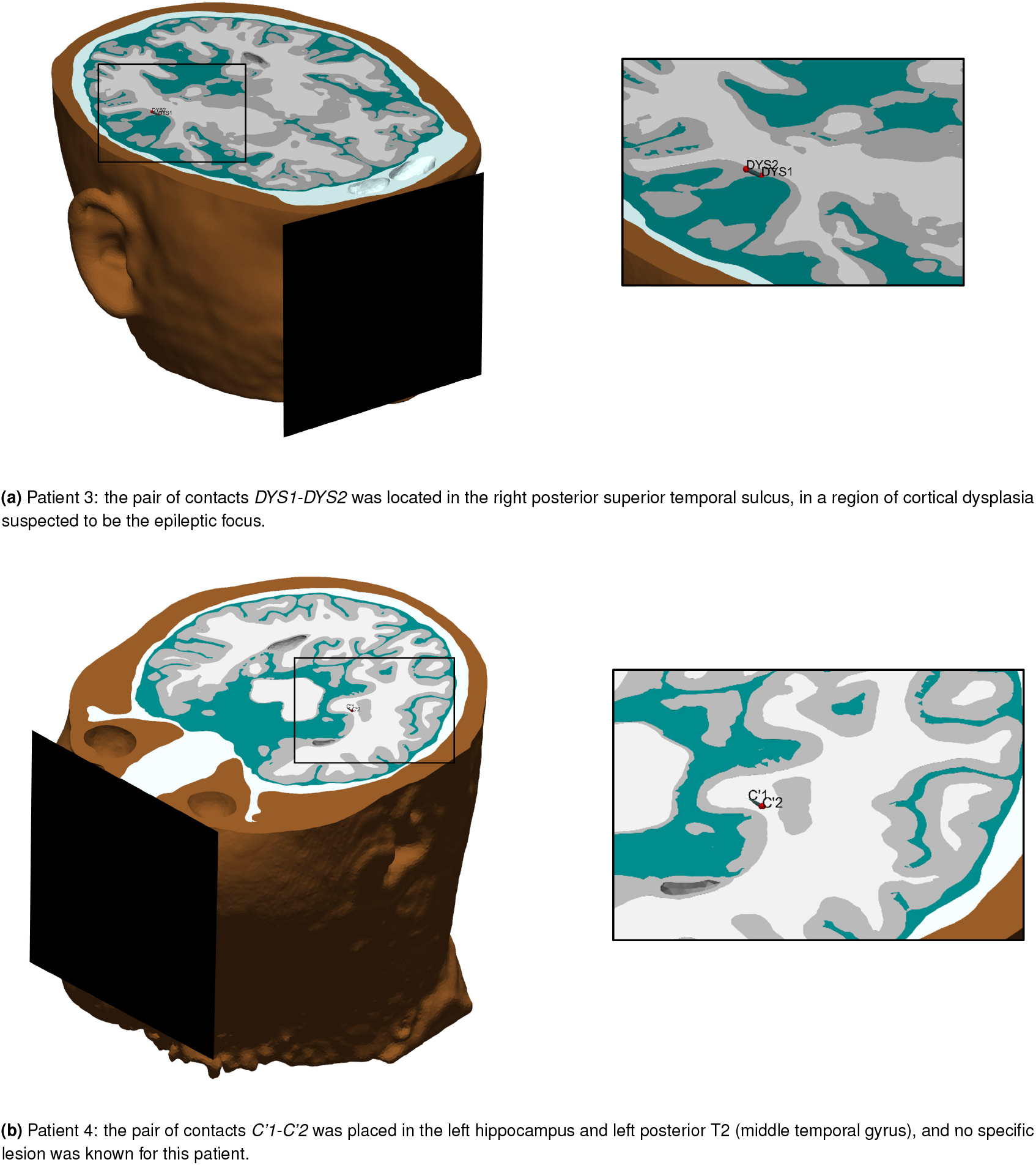
3D schematic view of the location of the most epileptogenic SEEG contacts for each patient (one pair in each case, since SEEG signals were obtained from bipolar derivations).

**Fig. S2.**
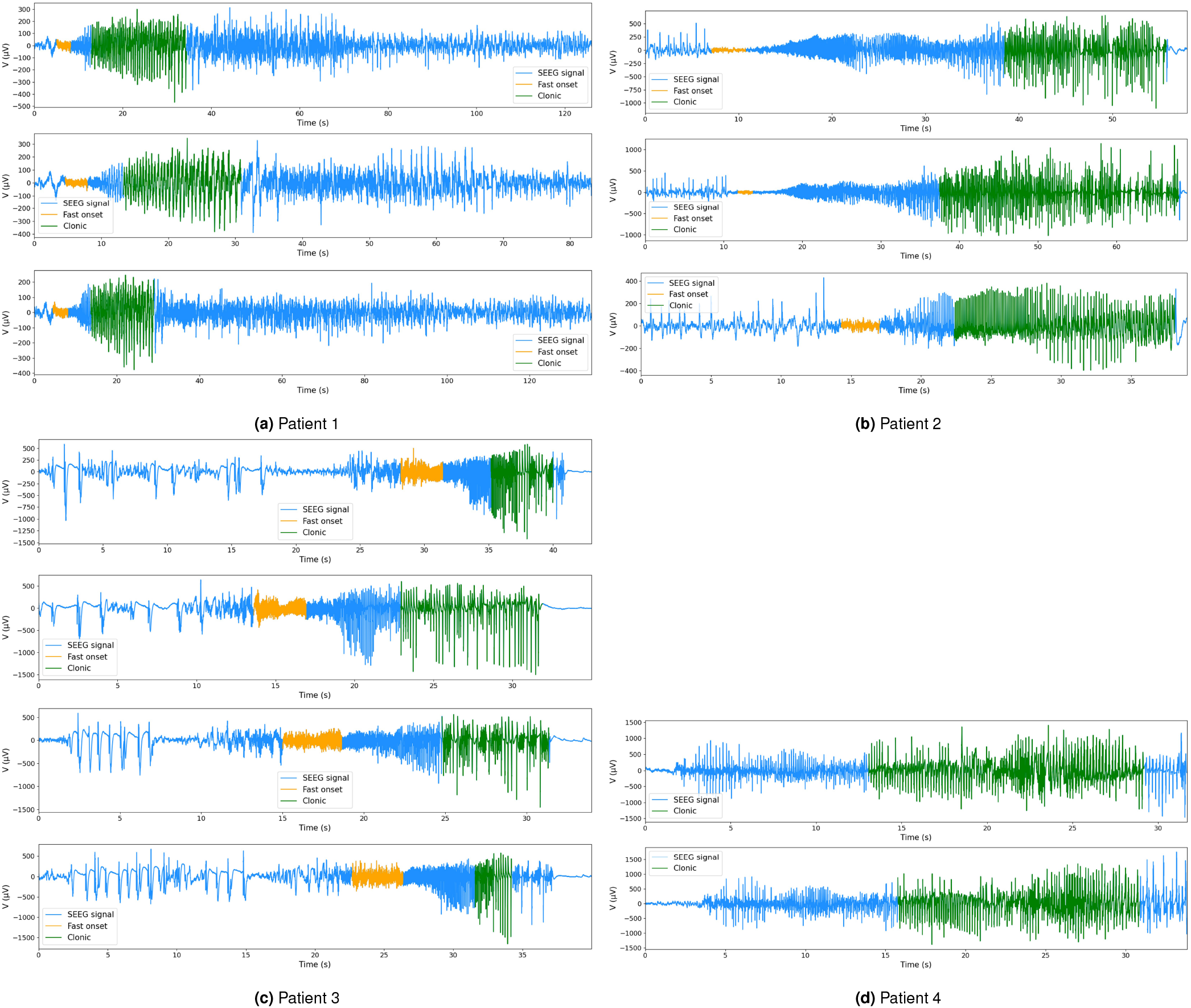
SEEG recordings during seizure events for all the patients included in the study (three seizures for Patient 1 and 2, four seizures for Patient 3 and two seizures for Patient 4). For each seizure, the periods selected for the calculation of the average representative frequency of the LFVA and clonic phases are shown in yellow and green respectively.

**Fig. S3.**
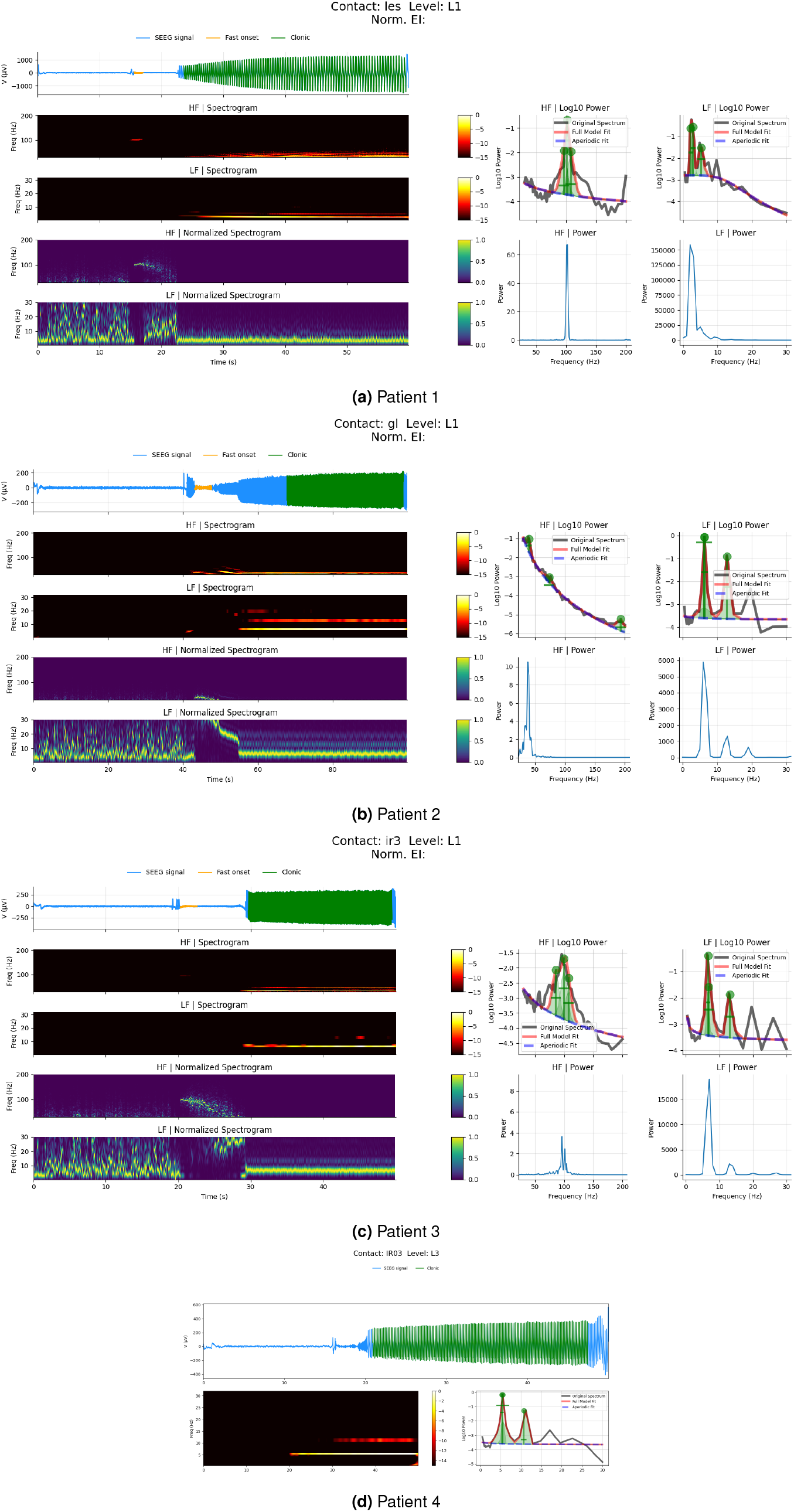
Simulated seizures for the four patients. Top panels display the simulated SEEG signals. The periods selected to compute the main frequency of each phase, reported in Table E1 are marked in yellow (fast onset) and green (rhythmic ictal period). The high- and low-frequency spectrogram and normalized spectrogram (see Appendix A2), as well as the power spectral density (PSD) as also shown.

**Fig. S4.**
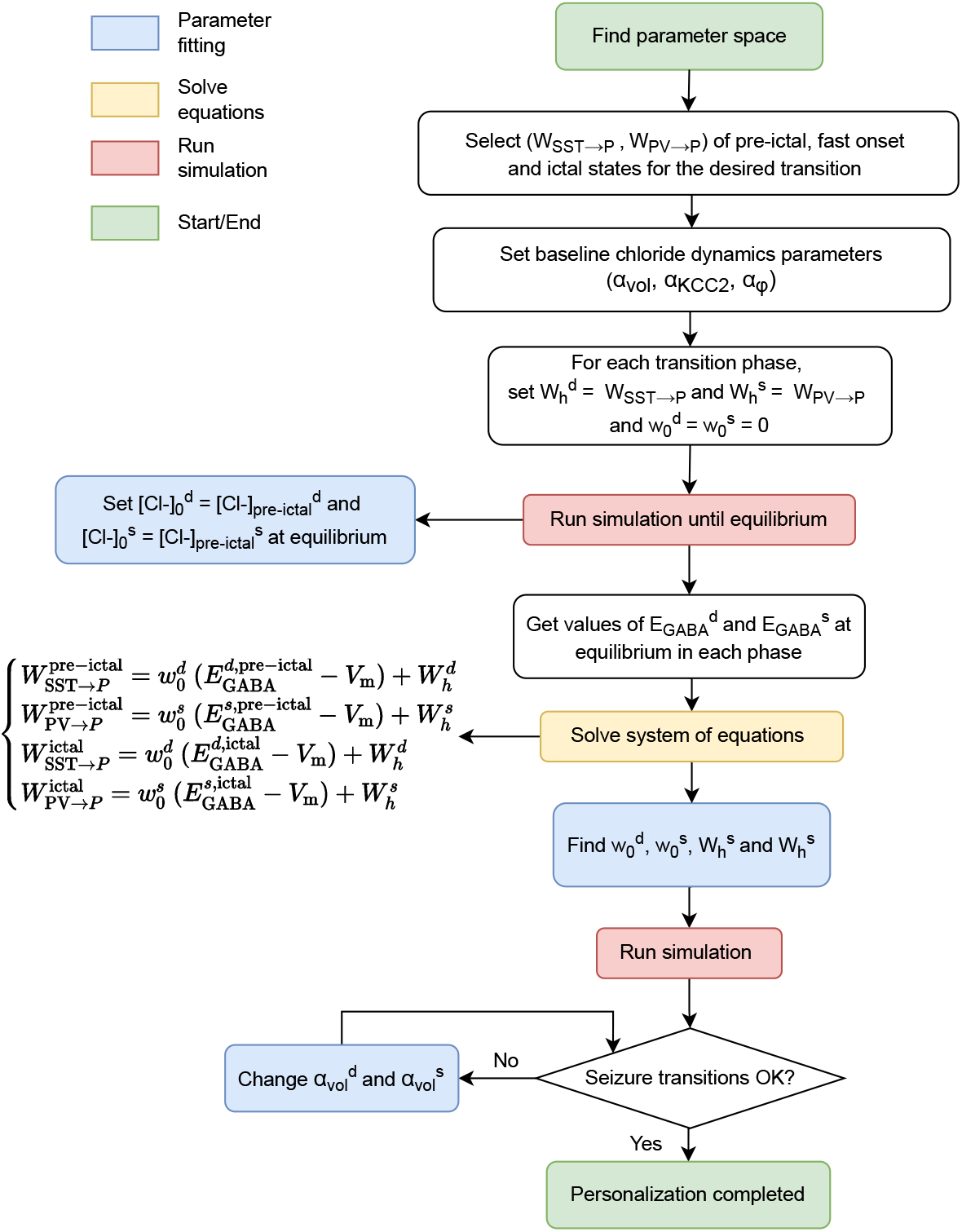
Flowchart summarizing the process for personalization of chloride dynamics parameters (see Table 2 in the main text) based on patient’s SEEG data. Starting from the personalized parameter space for the patient (Figure 2 in the main text), we must find the set of chloride accumulation parameters that will drive the system through the desired transition phases from interictal to seizure, reproducing the transition observed in the patient’s SEEG data.

## Footnotes

‡ Of note, our NMM includes a synapse not present in the “Wendling-class” model presented in [1], the autaptic connection in the PV cell populations. Our baseline parameters for this synapse are: *W*_PV→PV_ = −10 mV, *C*_PV→PV_ = 300, 1*/τ*_PV→PV_ = 500 s^−1^.

